# Heat Shock Factor 1 Drives Intergenic Association of Its Target Gene Loci Upon Heat Shock

**DOI:** 10.1101/400499

**Authors:** Surabhi Chowdhary, Amoldeep S. Kainth, David Pincus, David S. Gross

**Author notes:** Equal Contribution (Surabhi Chowdhary, Amoldeep S. Kainth).

## Abstract

Transcriptional induction of *Heat Shock Protein* (*HSP*) genes in yeast is accompanied by dynamic changes in their 3D structure and spatial organization, yet the molecular basis for these phenomena remains unknown. Using chromosome conformation capture and single cell imaging, we show that genes transcriptionally activated by Heat Shock Factor 1 (Hsf1) specifically interact across chromosomes and coalesce into diffraction-limited intranuclear foci. Genes activated by the alternative stress regulators Msn2 and Msn4, in contrast, do not interact among themselves nor with Hsf1 targets. Likewise, constitutively expressed genes, even those interposed between *HSP* genes, show no detectable interaction. Hsf1 forms discrete subnuclear puncta when stress-activated, and these puncta dissolve in concert with transcriptional attenuation, paralleling the kinetics of *HSP* gene coalescence and dissolution. Nuclear Hsf1 and RNA Pol II are both necessary for intergenic *HSP* gene interactions, while DNA-bound Hsf1 is necessary and sufficient to drive coalescence of a heterologous gene. Our findings demonstrate that Hsf1 can dynamically restructure the yeast genome.

## INTRODUCTION

Increasing evidence suggests that nuclear processes such as transcription, recombination and repair can be influenced not only by local chromatin structure but also by three-dimensional genome architecture. Genomes of higher eukaryotes are compartmentalized into discrete structural and regulatory units termed topologically associating domains (TADs) (reviewed in Dekker and Mirny, 2016; Hnisz et al., 2016a; Wendt and Grosveld, 2014). Genes located within these structures tend to have similar expression states and epigenetic signatures, and perturbation of TAD integrity may lead to aberrant activation (Flavahan et al., 2016; Hnisz et al., 2016b; Lupianez et al., 2015). Recent studies using genome-wide, chromosome conformation capture (3C)-based techniques have unveiled spatial genomic structures analogous to mammalian TADs in budding yeast (Eser et al., 2017; Hsieh et al., 2015).

DNA looping has been directly implicated in transcriptional control. For example, the DNA between enhancers and cognate promoters, typically 20-50 kb in mammals, loops out, permitting physical contact between these regulatory regions (reviewed in Furlong and Levine, 2018). Analogous, albeit smaller, enhancer (UAS)-promoter loops have been observed in *S. cerevisiae* (Chowdhary et al., 2017; Dobi and Winston, 2007). Physical contacts between the 5’ and 3’ ends of actively transcribed genes, as well as between gene regulatory elements and actively transcribed coding sequences, have been observed in both yeast (Chowdhary et al., 2017; Hampsey et al., 2011) and mammals (Beagrie et al., 2017; Lee et al., 2015).

Importantly, DNA loops tend to be dynamic, and such dynamism facilitates long-range chromosomal interactions. For example, both proximally and distally located mammalian genes engage in frequent contacts that may contribute to their co-regulation (Fanucchi et al., 2013; Li et al., 2012; Rao et al., 2014; Zhang et al., 2013). Moreover, activated mammalian genes have been observed to reposition themselves into discrete sites of intense RNA synthesis termed “transcription factories” (Osborne et al., 2004; Papantonis et al., 2012; Papantonis et al., 2010; Park et al., 2014; Schoenfelder et al., 2010). In these and other examples, it is thought that increased transcription is fostered by high local concentrations of RNA Pol II and pre-mRNA processing factors present in pre-existing stable substructures (Feuerborn and Cook, 2015; Mitchell and Fraser, 2008). However, a single molecule analysis using super-resolution microscopy indicated that Pol II clusters form transiently, with their mean lifetime increasing upon transcriptionally stimulating conditions (Cisse et al., 2013). Therefore, whether all transcriptionally active genes cluster, whether such clustering is the cause or consequence of transcription, what factors mediate clustering, and to what extent this mode of transcriptional control exists in eukaryotes other than mammals remains unknown.

A powerful model with which to study dynamic genome restructuring is the heat shock (HS)-responsive family of genes in *S. cerevisiae*. Many of these genes, including those encoding molecular chaperones and cytoprotective heat shock proteins (HSPs), are under the regulation of Heat Shock Factor 1 (Hsf1), an evolutionarily conserved, gene-specific activator (Gomez-Pastor et al., 2018; Wu, 1995). Genes under the regulation of Hsf1 undergo dramatic transformations in chromatin structure upon their activation. These alterations include gene-wide disassembly of nucleosomes (Zhao et al., 2005) and substantial increases in Hsf1, Mediator, SAGA and Pol II occupancy (Fan et al., 2006; Kim and Gross, 2013; Kremer and Gross, 2009; Vinayachandran et al., 2018). Additionally, prominent intragenic and intergenic chromosomal contacts accompany *HSP* gene activation. These alterations include DNA looping between UAS and promoter, promoter and terminator, as well as between regulatory regions and coding sequences. Activated *HSP* genes also engage in frequent *cis-* and *trans-*interactions with each other, dynamically coalescing into diffraction-limited foci (Chowdhary et al., 2017). It is unknown what underlies these genomic rearrangements. It is also unclear whether gene clustering is the default state for transcriptional control in budding yeast, as suggested for mammalian cells.

Here we demonstrate that stress-activated Hsf1 is a key determinant driving interactions between yeast *HSP* gene loci during heat shock. Thermal stress-responsive genes activated by alternative activators – Msn2 and Msn4 – do not detectably cluster, nor do coordinately regulated ribosomal protein genes. While high levels of transcription are necessary for coalescence, they are not sufficient. Our results argue against the idea that gene repositioning is a general feature of transcriptional activation and instead point to activators such as Hsf1 as the drivers of global genome restructuring.

## RESULTS

### *HSP* Genes Engage in Robust Intergenic Interactions That Exclude Genes Interposed Between Them

*HSP* genes engage in extensive intra-and interchromosomal interactions upon their heat shock-induced activation (Chowdhary et al., 2017). If *HSP* gene coalescence is biologically significant, then one might predict that non-*HSP* genes would be excluded from such clustering, even those residing in close linear proximity. To test this, we used a modified version of 3C to investigate intergenic interactions within a 35 kb domain on Chr. XII (TaqI-3C; see Materials and Methods and ***Reviewer’s Appendix***). Three *HSP* genes – *UBI4*, *HSP104* and *SSA2 –* lie within this domain and the promoter of each is inducibly occupied by Hsf1 in cells exposed to acute heat shock (30° to 39°C shift for 5 min; Figure 1A). Under non-heat shock (NHS) conditions, no 3C interactions above background could be detected between these genes (Figure 1B (blue matrices); gene regions defined in Figure S1), consistent with their low basal transcription and previous nucleosome-resolution chromatin contact analysis indicating that these genes lie within separate chromosome interaction domains (CIDs) (Hsieh et al., 2015).

**Figure 1.**
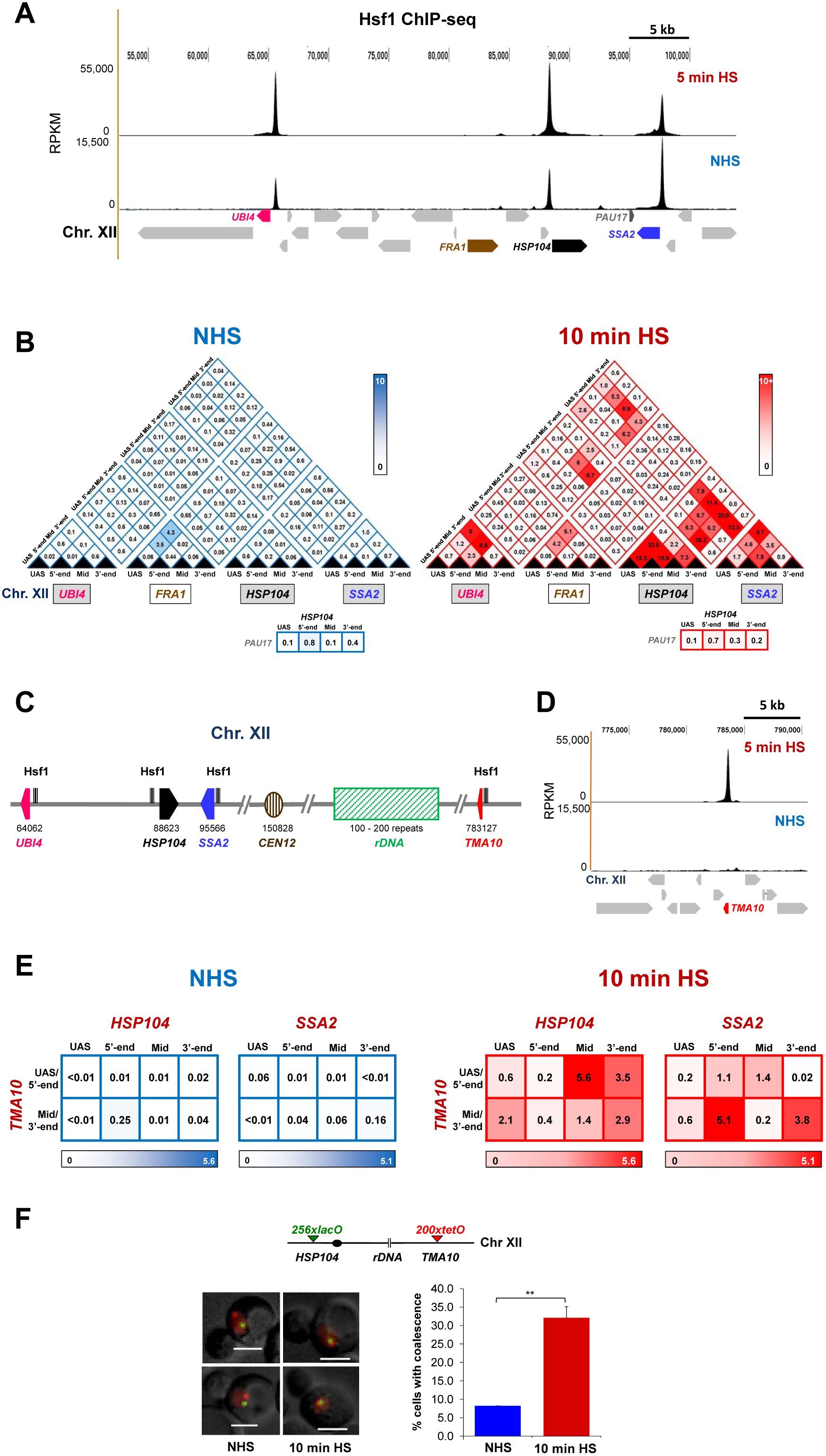
Hsf1-Regulated Genes Engage in Highly Specific Intergenic Interactions. (A) Hsf1 ChIP-seq profile of a 50 kb region on the left arm of Chr. XII in NHS and 5 min HS cells. Genes subjected to TaqI-3C analysis in panel B are highlighted. RPKM, reads per kilobase per million mapped reads.
(B) (*Left*) Contact frequencies between the indicated regions of *UBI4, FRA1, HSP104* and *SSA2* in haploid cells (strain BY4741) grown under NHS conditions (30°C), as determined by TaqI-3C. Values indicate normalized interaction frequencies, determined as described in Materials and Methods. Those ≤0.1 are indistinguishable from background. Gene regions are defined in Figures S1A and S1B. (*Right*) Same, except chromatin was isolated from cells exposed to a 10 min, 39°C heat shock. *HSP104* - *PAU17* contact frequencies are shown below triangulated analysis. Intensity of color is proportional to the frequency of interaction. Data are derived from two independent biological replicates (qPCR=4 for each primer combination).
(C) Physical map of Chr. XII. The rDNA repeats and *TMA10* are located on the right arm of Chr. XII, 600 kb and 300 kb from *TEL12R*, respectively. Coordinates are numbered relative to the left telomere and do not take into account the presence of rDNA.
(D) Hsf1 occupancy profile of the indicated ~20 kb region on the right arm of Chr. XII. Hsf1 ChIP-seq analysis presented as in A.
(E) Intergenic interaction frequencies between *HSP104-TMA10* and *SSA2-TMA10* in NHS and 10 min HS cells. For each pairwise test, N=2, qPCR=4.
(F) Fixed-cell fluorescence microscopy of diploid strain ASK727 (*HSP104-lacO_256_ TMA10-tetO_200_ GFP-LacI TetR-mCherry*). *(Top)* Schematic of *lacO*-tagged *HSP104* and *tetO*-tagged *TMA10* loci (filled circle, centromere). (*Bottom left*) Micrographs of representative cells under NHS and 10 min HS conditions. Scale bar = 2 μm. (*Bottom right*) Percentage of cells exhibiting coalescence. 50-80 cells were evaluated per sample at each time point. N = 2; **, *P*< 0.01 (calculated using one-tailed t-test).

However, following heat shock, not only did neighboring *HSP104* and *SSA2* engage in intense interactions, but *UBI4* frequently contacted both genes (Figure 1B (red matrices)). This is despite the fact that *UBI4* is separated from *HSP104* by 25 kb and from *SSA2* by 33 kb, distances encompassing six and eight CIDs, respectively (Figure S2A). By contrast, the constitutively active and looped gene, *FRA1*, located between *UBI4* and *HSP104*, engaged in no detectable interactions with *HSP104, UBI4* or *SSA2* under either condition (Figure 1B). Likewise, *PAU17*, a non-*HSP* gene interposed between *HSP104* and *SSA2*, failed to engage in physical interactions with either gene. Thus, both *FRA1* and *PAU17*, despite being transcriptionally active and residing in close proximity to the *HSP* genes, are excluded from the heat shock-mediated intergenic interactions taking place around them.

We next wished to know if *cis*-intergenic interactions are spatially confined by chromosomal location. Towards this end, we asked whether *HSP104* and *SSA2*, residing on the left arm of Chr. XII, would physically interact with *TMA10*, a gene located on the distal right arm and inducibly occupied by Hsf1 (Figures 1C, 1D). Genome-wide 3C-based analyses have indicated that the left and right ends of Chr. XII are physically isolated from each other, due to a “near absolute” barrier conferred by the 100-200 rDNA repeats that assemble into the nucleolus (Cournac et al., 2012; Duan et al., 2010; Rutledge et al., 2015). Consistent with these prior studies, TaqI-3C failed to detect above-background interaction between *HSP104 - TMA10* or *SSA2 - TMA10* in non-induced cells (Figure 1E, *left*). However, following a 10 min heat shock, physical interactions between these Hsf1 targets were readily detectable (Figure 1E, *right*).

To validate this result using an orthogonal approach, we performed single-cell microscopy analysis of a heterozygous diploid bearing chromosomally linked *lacO*-tagged *HSP104* and *tetO*-tagged *TMA10* genes and co-expressing GFP-LacI and TetR-mCherry (Figure 1F). Under NHS conditions, the two genes rarely co-localized (in <10% of cells), a frequency that may arise from coincidental overlap. However, following 10 min HS, the frequency of *HSP104-lacO_256_ - TMA10-tetO_200_* co-localization increased to 35%, a significant increase (*P*<0.01, one-tailed t-test), consistent with the above 3C analysis as well as imaging experiments below (Figures 3A and 3B). While it is possible that these intrachromosomal interactions are facilitated by loss of nucleolar integrity, fluorescence imaging of acutely heat-shocked cells expressing an RFP-tagged nucleolar protein argues otherwise (Figure S2B). These results demonstrate that *HSP* gene interactions are not only specific, but sufficiently robust to circumvent the physical barrier imposed by the nucleolus.

### Heat Shock-Induced Intergenic Interactions are Distinct to Hsf1 Target Genes

While a large number of genes are transcriptionally induced by heat shock, only a small fraction are dependent on Hsf1, as revealed by transcriptome-wide sequencing (RNA-seq) of Hsf1^+^ *vs.* Hsf1^−^ cells (Figure 2A) (Hsf1^−^ cells conditionally depleted of nuclear Hsf1; see below). Therefore, are intergenic interactions a general feature of heat-shock-activated genes or are they distinctive to Hsf1-regulated genes? Of particular interest are genes whose thermal-responsive regulation is under the control of Msn2 (and its paralogue, Msn4). Msn2/Msn4 (hereafter referred to as Msn2/4) regulates the transcription of 200-300 genes in response to a variety of environmental stresses, including heat, oxidative, osmotic and salt (Elfving et al., 2014; Gasch et al., 2000). We selected three genes – *CTT1*, *PGM2* and *RTN2 –* whose thermal stress-dependent transcription is dependent on Msn2/4 and independent of Hsf1 (Figures 2A, 2B, S3A) (Pincus et al., 2018).

**Figure 2.**
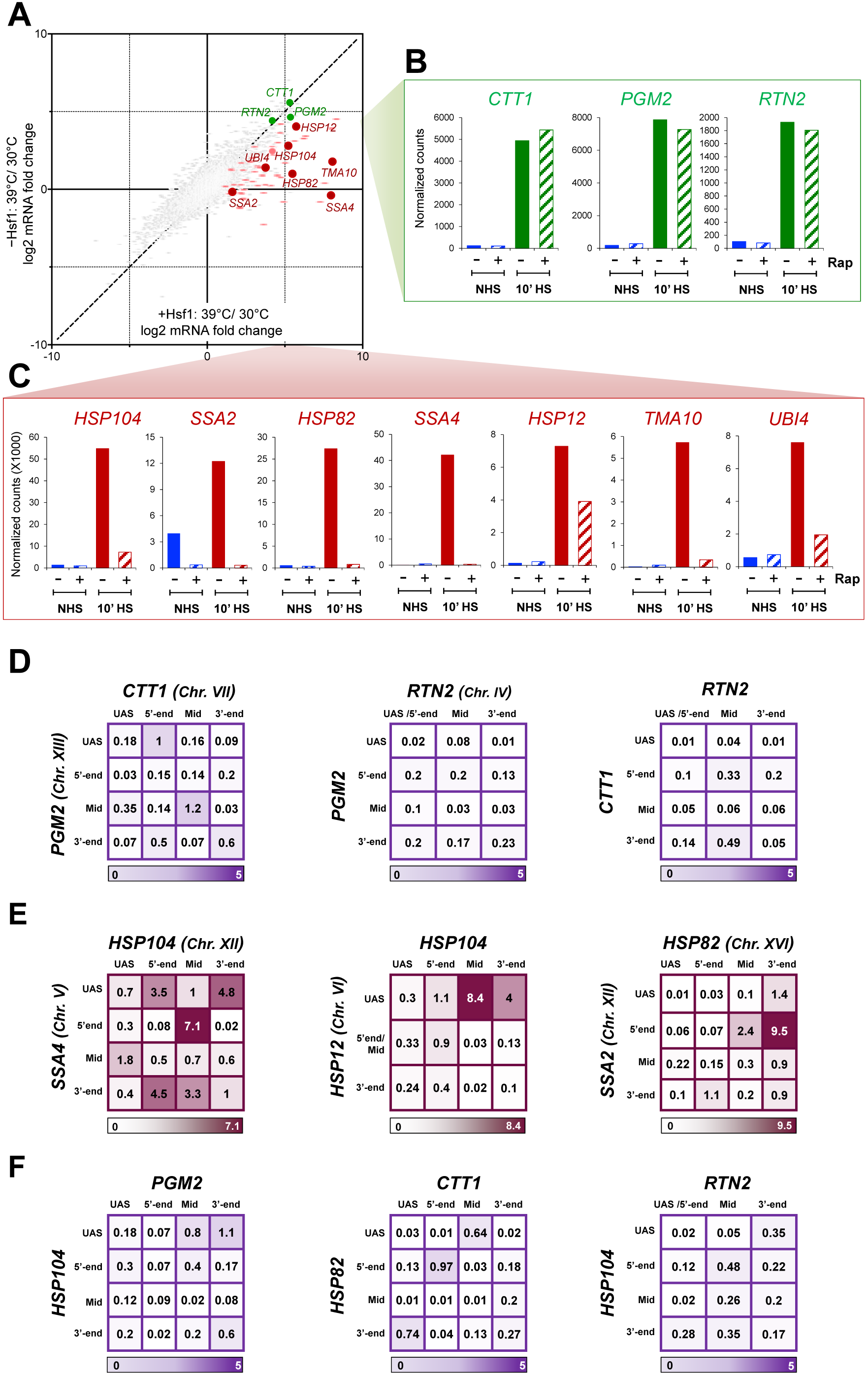
Msn2/4-Regulated Genes Are Strongly Induced by Heat Shock Yet Fail to Engage in Intergenic Interactions. (A) Transcriptome-wide HS/NHS fold change in expression in the presence and absence of nuclear Hsf1 using the Hsf1 Anchor Away system (Hsf1-AA). 1 μM rapamycin was added for 45 min to deplete Hsf1 from the nucleus (-Hsf1) or not (+Hsf1), prior to continued incubation at 30°C (NHS) or heat shock (HS) at 39°C for 10 min. Plotted are RNA-seq count ratios determined for every gene. Categories: Red/Pink, Hsf1 occupied genes in acutely heat-shocked cells (Pincus et al., 2018); Green, Msn2/4-dependent, Hsf1-independent genes (as defined in text); Grey, all others.
(B) Transcript levels (normalized RNA-seq reads) of select Msn2/4-dependent genes under each of the indicated conditions.
(C) As in B, except for select Hsf1-dependent genes.
(D) Matrix summaries of intergenic interaction frequencies between the indicated Msn2/4-target genes in 10 min HS cells (analysis and presentation as in Figure 1B). For each pairwise test, N=2, qPCR=4.
(E) As in D, except pairwise tests were conducted between the indicated Hsf1-target genes.
(F) As in D, except intergenic interactions between Msn2/4-and Hsf1-regulated genes were determined.

TaqI-3C analysis revealed the presence of heat-shock-dependent intragenic looping interactions within all three Msn2/4-target genes (Figure S3B; gene maps in Figure S1C), consistent with the notion that such restructuring is characteristic of actively transcribed genes (Chowdhary et al., 2017). Nonetheless, using primers corresponding to the UAS, 5’-end, mid-ORF and 3’-end of each gene, we were unable to detect above-background interactions between *CTT1*, *PGM2* and *RTN2* upon their transcriptional activation (Figures 2D, S3C) despite the presence of readily detectable interactions between Hsf1 target genes (Figures 2E, S3C) in the same 10 min heat-shocked cells. Moreover, we did not detect any interactions between heat shock-induced Msn2/4-target genes and Hsf1 target genes (Figures 2F and S3C).

We next asked if constitutively active genes associate with each other, and tested intergenic interactions between two coordinately regulated ribosomal protein genes, *RPL10* and *RPL22A*, as well as between two unrelated genes, *FAS2* and *RPL10*. Each gene is heavily transcribed under NHS conditions, as determined by both RNA-seq and nascent RNA measurements (Figure S4A; (Pincus et al., 2018)), yet no above-background interactions could be detected (Figure S4B). Together, these data suggest that neither heat shock-inducible Msn2/4-regulated genes nor other highly expressed, coordinately regulated genes interact either among themselves or with Hsf1 target genes.

Single cell imaging corroborated these findings. We observed that *HSP104-lacO_256_* and *HSP12-lacO_128_*, residing on Chr. XII and Chr. VI, respectively, coalesced into single diffraction-limited foci in >30% of cells under acute heat shock conditions (2.5 or 10 min heat shock), and the frequency of such coalescence was significantly higher than in either the control or 30 min heat-shocked state (Figure 3A (*solid bars)*, *P* < 0.05, one-tailed t-test). This is in agreement with our previous findings that *HSP* gene interactions are highly dynamic, detectable within 60 sec of heat shock, yet evanescent (Chowdhary et al., 2017). In contrast, only background levels of coalescence were observed between Hsf1-regulated *HSP104* and Msn2-regulated *PGM2* (Figure 3A, *striped bars*). Moreover, while the distance between *HSP104* and *HSP12* was normally distributed under NHS conditions, the distribution became skewed towards shorter distances during the acute stages of heat shock (Figure 3B), consistent with interchromosomal clustering of the two loci. No such change was observed between *HSP104* and *PGM2* in identically treated cells. Collectively, 3C and microscopy analyses suggest that (i) Msn2/4-regulated genes do not coalesce – either with themselves or with Hsf1-targets – in response to heat shock; and (ii) Hsf1 targets do not coalesce with other transcriptionally active genes, even those induced by thermal stress. These observations argue that coalescence may be a distinguishing feature of Hsf1-activated genes.

**Figure 3.**
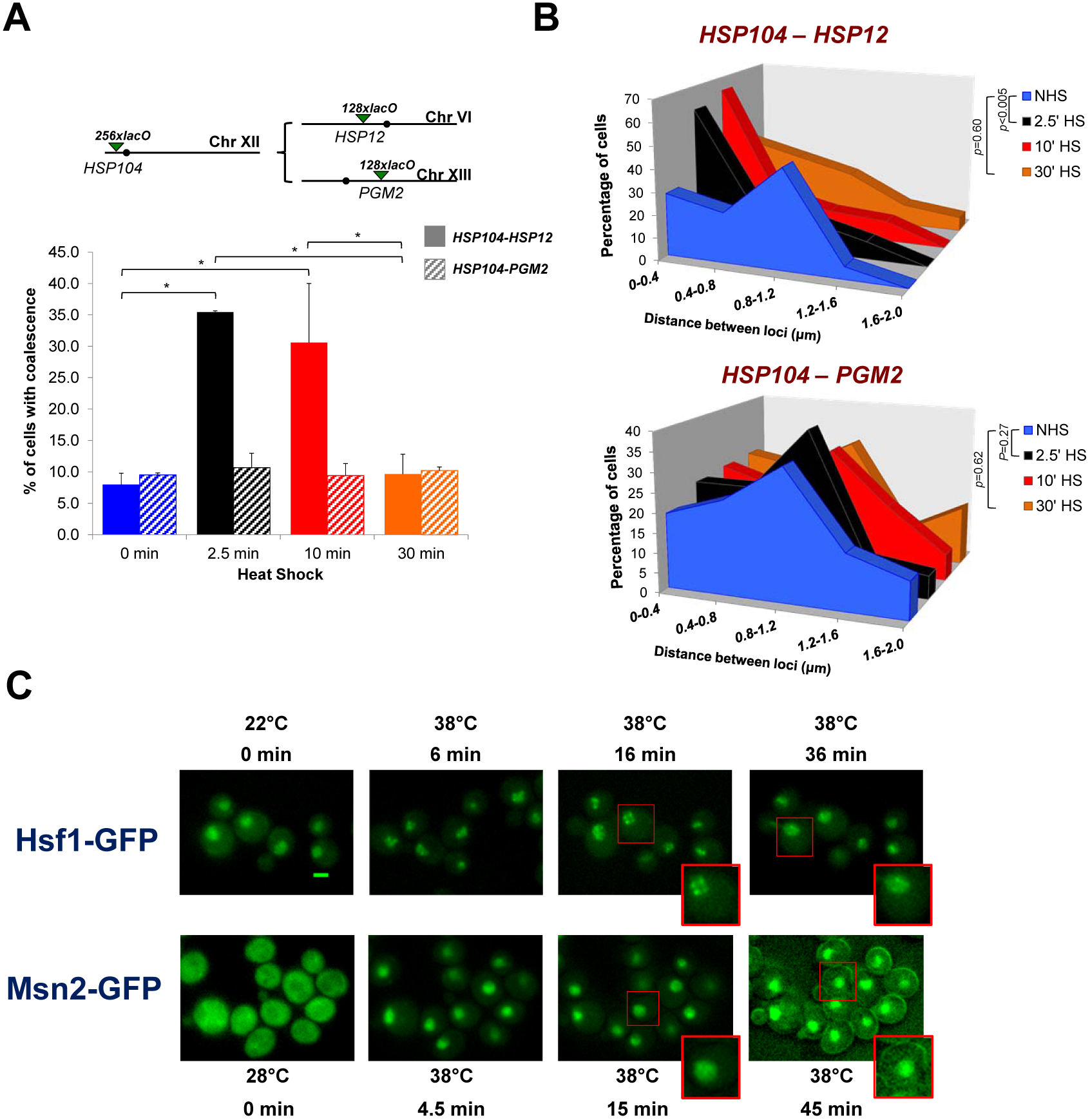
Single Cell Analysis Reveals Preferential, Dynamic Coalescence Between Hsf1-Target Genes Paralleled by Hsf1 Puncta Formation. (A) *(Top)* Schematic of *lacO*-tagged loci present in *HSP104* and *HSP12* or *PGM2* heterozygous diploids (filled circles, centromeres). (*Bottom*) Percentage of cells bearing tagged *HSP104-HSP12* (solid bars) or *HSP104-PGM2* (shaded bars) exhibiting coalescence as determined by fixed-cell fluorescence microscopy. For *HSP104-HSP12*, 50-70 cells were evaluated per sample at each time point (N=2); for *HSP104-PMA2*, 50-100 cells were evaluated per sample at each time point (N=2). *, *P*< 0.05 (calculated by ANOVA followed by Tukey’s post hoc analysis). Data for *HSP104-HSP12* coalescence are from a previous study (Chowdhary et al., 2017).
(B) Distribution of distances between *HSP104* and *HSP12* (*Top*) or *HSP104* and *PGM2* (*Bottom*) in cells subjected to the indicated conditions. Depicted is the percentage of cells with tagged loci within the indicated distance (binned at intervals of 0.4 μm) in a given population of fixed cells. This analysis encompasses a subset of cells analyzed in A, as only those cells where two spots lie in same plane can be taken into account. Distance across multiple planes could not be accurately measured due to lower resolution in the *Z* direction. *P* values were calculated by Wilcoxon rank sum test.
(C) Fluorescence microscopy of live cells expressing Hsf1-GFP (*Top*) or Msn2-GFP (*Bottom*) prior to or following application of heat for the times and temperatures indicated. Images are presented at same magnification; cells boxed in red are enlarged at bottom of the respective images. Scale bar = 2 μm.

### Hsf1 Forms Discrete Intranuclear Puncta in Cells Exposed to Thermal Stress

To address the possibility that Hsf1 itself coalesces upon activation, we imaged live cells harboring Hsf1-GFP. As shown in Figure 3C, Hsf1 is largely nuclear and diffusely localized under NHS conditions. Following brief heat shock (6 − 16 min), the protein forms discrete nuclear puncta. However, by later time points (36 − 66 min), Hsf1 puncta dissolve, and the distribution of intranuclear Hsf1 returns to a diffuse state, closely paralleling the kinetics of *HSP* gene coalescence and dissociation described above. If the formation of Hsf1 puncta reflects coalescence of its gene targets, then it might be predicted that Msn2, despite strongly activating transcription in response to heat shock (Figures 2B and S3A) will not form nuclear puncta. As shown in Figure 3C, Msn2-GFP, largely cytoplasmic in NHS cells (0 min), translocates to the nucleus under acutely stressful conditions (4.5 min), consistent with previous reports (Chi et al., 2001; Gorner et al., 1998). In contrast to Hsf1, the intranuclear distribution of Msn2 remains diffuse throughout the heat shock time course (Figure 3C, 4.5 − 45 min). Notably, exposure to 8.5% ethanol has similar effects, including the transient formation of Hsf1 puncta (L.S. Rubio, A.S.K. and D.S.G., unpubl. observations), so the ability/inability to form nuclear puncta may be an inherent property of Hsf1 and Msn2 activation.

### Hsf1 and Pol II are Necessary to Drive Interactions Between *HSP* Genes During Heat Shock

To directly test the importance of Hsf1 in driving changes in *HSP* gene conformation and 3D nuclear organization, we conditionally depleted it from the nucleus using the Anchor Away technique (Haruki et al., 2008). Growth of *HSF1-FRB* cells on rapamycin demonstrates that Hsf1 is essential for viability, even at 30°C (Figure 4A), consistent with previous observations (Sorger and Pelham, 1988). Cytoplasmic sequestration of Hsf1-FRB, achieved by pre-exposure of cells to rapamycin for 90 min, depleted Hsf1-FRB occupancy of representative *HSP* genes following a subsequent 10 min heat shock (e.g., Figure 4B, *left*). Consistent with Hsf1’s central role in regulating these genes (Pincus et al., 2018; Solis et al., 2016), Pol II occupancy was likewise severely reduced (Figure 4B (*right*)), as was transcript accumulation (Figure 2C). Concomitant with this reduction in transcription, formation of 5’-3’ gene loops was obviated, as were other intragenic interactions including UAS - promoter looping (Figure 4C, pink bars). In conjunction, intergenic coalescence was reduced to background levels (Figure 4D). As expected, intragenic looping of constitutively expressed *BUD3* was unaffected by this perturbation (Figure S5A). Therefore, Hsf1 is required to drive its target genes into a looped and physically interactive state in response to heat shock.

**Figure 4.**
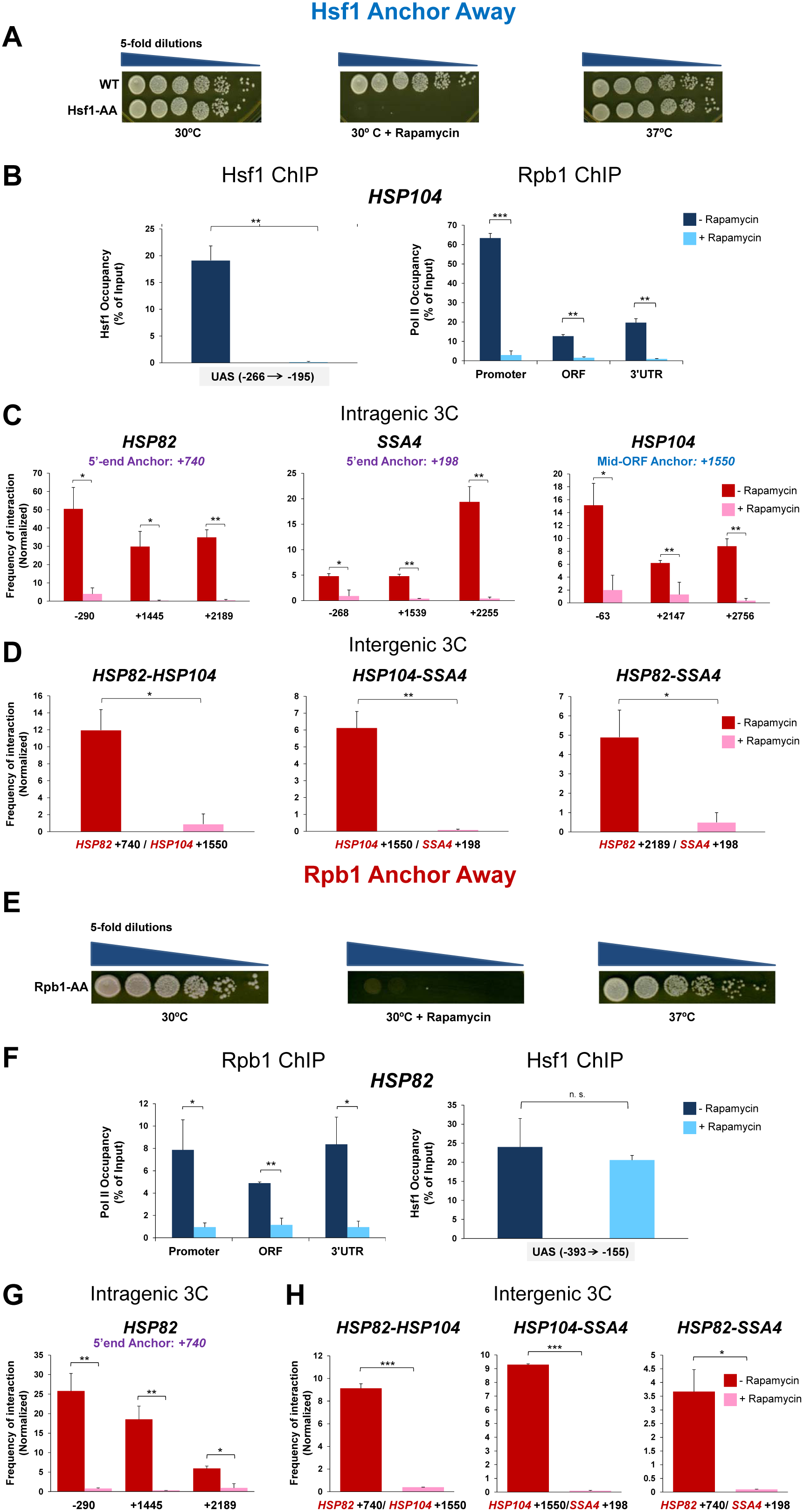
Nuclear Hsf1 and Pol II Are Necessary for Driving Intergenic *HSP* Gene Interactions. **(A)** Spot dilution analysis of Hsf1-AA (BY4742-HSF1-AA) cells. Five-fold serial dilutions of cells were spotted onto YPDA +/- rapamycin as indicated. Plates were incubated at 30° or 37°C for 2-3 days. WT, BY4742. **(B)** Hsf1-FRB-yEGFP and Rpb1 ChIP analysis of *HSP104*. Hsf1-AA cells were pretreated with rapamycin for 90 min or not as indicated, then subjected to a 10 min HS and processed for ChIP. Depicted are means + SD (N=2; qPCR=4). **, *P*<0.01; ***, *P*<0.001 (calculated using one-tail t-test). **(C and D)** TaqI-3C analysis of *HSP* intragenic and intergenic interactions in Hsf1-AA cells pretreated with rapamycin followed by 10 min heat shock. Depicted are means + SD (N=2; qPCR=4). *, *P*<0.05; **, *P*<0.01 (calculated as in B). **(E)** Spot dilution analysis of Rpb1-AA cells (yFR1324) conducted as described in A. **(F)** Rpb1-FRB and Hsf1 ChIP analysis of *HSP82.* Rpb1-AA cells were pretreated with rapamycin for 60 min or not as indicated, followed by a 10 min HS. Depicted are means + SD (N=2; qPCR=4). *, *P*<0.05; **, *P*<0.01; n.s. (not significant), *P*>0.05 (as in B). **(G and H)** Taq I − 3C analysis of *HSP* intragenic and intergenic interactions in Rpb1-AA cells pretreated with rapamycin or not (as indicated) followed by a 10 min HS. Depicted are means + SD (N=2; qPCR=4). *, *P*<0.05; **, *P*<0.01; ***, *P*<0.001 (as in B).

We next asked whether Pol II, in particular its largest subunit (Rpb1), is likewise required for the changes observed in *HSP* gene conformation and nuclear organization. Rapamycin-induced cytoplasmic sequestration of Rpb1-FRB rendered cells inviable on solid medium (Figure 4E). In cells pre-exposed to rapamycin for 60 min, Rpb1-FRB occupancy of heat shock-induced *HSP* gene promoters and coding regions was greatly reduced, although this nuclear depletion had little or no effect on Hsf1 occupancy (e.g., Figure 4F). Nonetheless, all intragenic interactions tested were greatly diminished by prior removal of Rpb1, including UAS-promotor looping (Figure 4G, first pairwise test), implicating Pol II in the stable formation of such loops. Concomitant with loss of intragenic looping was loss of all tested intergenic interactions (Figure 4H, pink bars), implicating Pol II and/or transcription in the interactions among *HSP* genes. As expected, intragenic looping interactions at *BUD3* were also diminished by Rpb1 depletion (Figure S5B). Therefore, Pol II is critical for the formation of novel intergenic interactions characteristic of activated *HSP* genes, yet even high levels of it – as inferred from intragenic looping, ChIP and expression assays – are not sufficient.

### Preventing Hsf1 DNA Binding Uncouples *HSP12* Coalescence from Transcription

Can intergenic *HSP* interactions be decoupled from transcription? To address this question, we analyzed the *HSP12* gene as a test case since it is regulated by both Hsf1 and Msn2/4. *HSP12* has an Hsf1 binding site (Heat Shock Element (HSE)) consisting of TTCn-NNNNN-nTTCn-NNNNN-nTTC, lying ~800 bp upstream of its transcription start site (TSS) and inducibly occupied by Hsf1 (Figure 5A). It also has seven stress-response elements (STREs) – CCCCY motifs – recognized by Msn2/4 located between the HSE and TSS. Conditional nuclear depletion of Hsf1 caused a moderate reduction in the heat shock-induced transcript level of *HSP12* as measured by RNA-seq (Figure 2C). Likewise, this perturbation caused a moderate reduction in heat shock-induced gene looping (Figure 5B). By contrast, Hsf1 depletion obviated *HSP12* interactions with *HSP104*, *HSP82* and *SSA2* altogether (Figure 5C). Thus, although *HSP12* transcription is only mildly affected by Hsf1 depletion, its interchromosomal interactions with other *HSP* genes are abolished. This observation suggests that *HSP* gene transcription, while strongly correlated with intragenic looping, can be uncoupled from intergenic coalescence.

**Figure 5.**
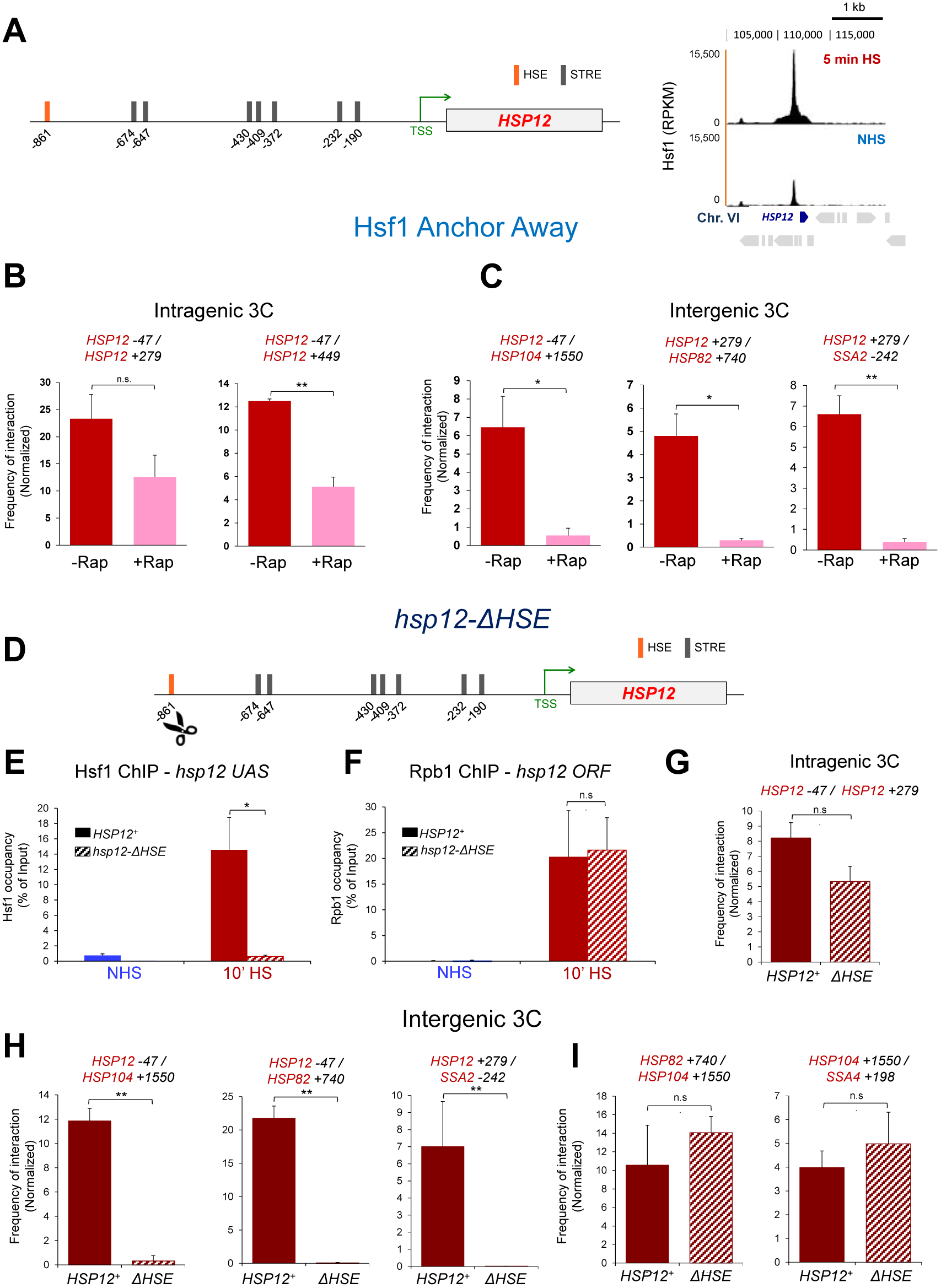
Hsf1 Depletion or Chromosomal HSE Deletion Uncouples *HSP12* Gene Transcription from Intergenic *HSP* Gene Interaction. (A) *(Left)* Physical map of *HSP12* depicting the HSE and STREs within its upstream region (derived from www.yeastract.com). Coordinates are numbered with respect to ATG. *(Right)* Hsf1 ChIP-seq profile of the indicated region on Chr. VI in NHS and 5 min HS cells.
(B) TaqI-3C analysis of *HSP12* intragenic interactions in Hsf1-AA cells subjected to a 10 min HS. Analysis and presentation as in Figure 4C. **, *P*<0.01; n.s., *P*>0.05.
(C) As in B, except frequency of intergenic interactions between *HSP12* and representative *HSP* genes was determined.
(D) Chromosomal DNA sequence encompassing a consensus HSE was excised to create an allele termed *hsp12-∆HSE* (see Materials and Methods).
(E) Hsf1 ChIP analysis was conducted on *HSP12^+^* and *hsp12-∆HSE* cells (strains AJ303 and AJ305, respectively) maintained at 30°C (NHS) or subjected to a 10 min HS as indicated. Analysis and presentation as in Figure 4B. *, *P*<0.05.
(F) Pol II (Rpb1) ChIP analysis conducted as in E.
(G) Intragenic contact frequencies of *HSP12* and *hsp12-∆HSE* following 10 min HS as deduced by TaqI-3C (N=2; qPCR=4). Presentation as in B.
(H) Intergenic contact frequencies between *HSP12* or *hsp12-∆HSE* and the indicated gene loci following 10 min HS. Presentation as in C.
(I) As in H, except intergenic contact frequencies between representative *HSP* genes in strains AJ303 and AJ305 were determined.

To further strengthen this point, we performed the complementary experiment by chromosomally excising the HSE from the genome upstream of *HSP12*, creating an allele termed *hsp12-∆HSE* (Figure 5D). This manipulation reduced Hsf1 occupancy to near-background levels under both NHS and 10 min HS conditions (Figure 5E). However, consistent with the Hsf1 Anchor Away results, there was little effect on either Pol II occupancy or intragenic looping (Figures 5F, 5G), in line with the notion that Hsf1 is largely dispensable for *HSP12* transcription under these conditions. Despite this, the HSE deletion suppressed *hsp12* interaction with *HSP104*, *HSP82* or *SSA2* (Figure 5H, compare solid vs. striped bars). As expected, other interchromosomal *HSP* gene interactions were unaffected (Figure 5I). These observations demonstrate that even when Hsf1 is largely dispensable for stimulating transcription, it is necessary for driving a target gene into physical interactions with other *HSP* genes.

### Ectopic Targeting of Hsf1 Is Sufficient to Drive Intergenic Association of a Heterologous Gene

Finally, to test whether DNA-bound Hsf1 is sufficient to cause an otherwise unrelated gene to coalesce with *HSP* genes, we chromosomally integrated a high-affinity Hsf1 binding site upstream of *BUD3*, creating an allele termed *UAS_HS_-BUD3* (Figures 6A and S1D). We then conducted TaqI-3C analysis on both non-induced and acutely heat-shock-induced cells. As shown in Figure 6B, Hsf1 strongly occupied *UAS_HS_-BUD3* but not *BUD3^+^* in response to a 10 min heat shock. Moreover, *UAS_HS_-BUD3* exhibited increased levels of intragenic looping interactions following heat shock (Figure 6C), consistent with its increased transcription (A.S.K. and D.S.G., unpubl. observations). Most significantly, *UAS_HS_-BUD3* engaged in novel intergenic interactions with both *HSP104* and *HSP82* (Figure 6D, *right*), in marked contrast to the wild-type gene (Figure 6D, *left*). Therefore, DNA-bound Hsf1 is sufficient to direct a heterologous gene into novel interactions with *HSP* genes in response to heat shock.

**Figure 6.**
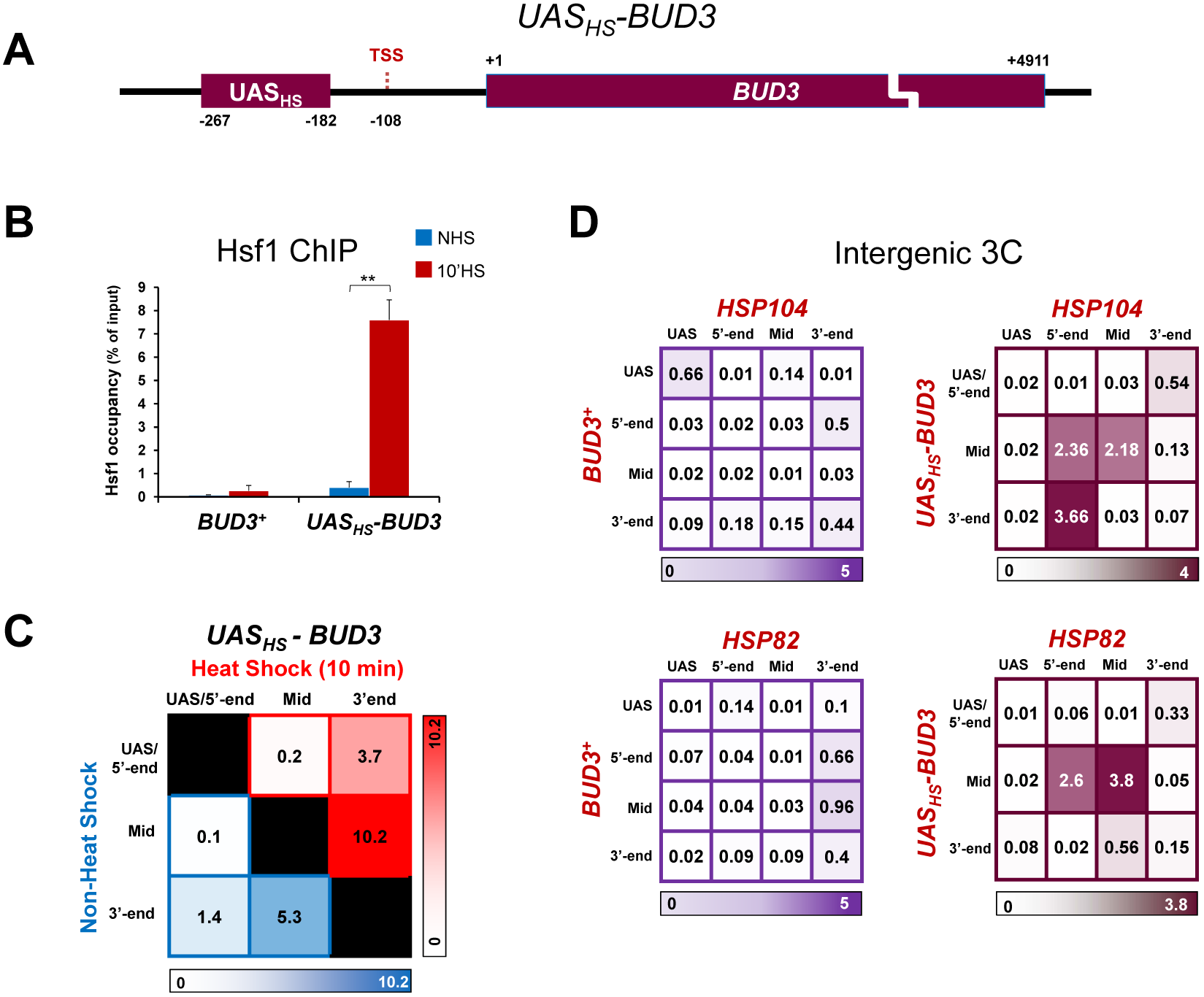
Ectopic, Heat Shock-Activated Hsf1 Is Sufficient to Drive Coalescence of a Heterologous Gene. (A) Physical map of the chromosomal transgene *UAS_HS_-BUD3*.
(B) Hsf1 occupancy of *BUD3* and *UAS_HS_-BUD3* (strains BY4741 and ASK804, respectively) under NHS and 10 min HS states. Analysis and presentation as in Figure 4B. **, *P*<0.01.
(C) Summary of intragenic interactions detected within *UAS_HS_-BUD3* in NHS and 10 min HS states (N=2; qPCR=4).
(D) Intergenic contact frequencies between *BUD3* or *UAS_HS_-BUD3* and the indicated *HSP* genes in 10 min HS cells (N=2; qPCR=4).

## DISCUSSION

### Hsf1 Target Genes Distinctively Interact Among Themselves Upon Activation

We present evidence using both molecular and single-cell imaging approaches that physical interactions between heat shock-activated Hsf1 target genes, located on the same or different chromosomes, are specific and robust. Genes interposed between Hsf1 targets are excluded from these interactions, yet *HSP* genes spatially segregated by rDNA repeats that comprise the nucleolus readily interact. Importantly, the *HSP* genes tested were located at a variety of chromosomal latitudes (Table S1), thereby rendering unlikely that the interchromosomal interactions we detect reflect fortuitous alignment of these genes along Rabl-organized chromosomes (Taddei and Gasser, 2012). Not all heavily transcribed genes coalesce, even those whose transcription is activated by alternative thermal stress-responsive activators. Likewise, coordinately regulated ribosomal protein genes show no detectable interaction despite the fact that the genes tested lie on the same chromosome and are separated by only 20 kb. The latter observation is consistent with genome-wide 3C analyses that failed to uncover significant interactions between Pol II genes across the yeast genome under the control condition (NHS) of our experiments (Duan et al., 2010; Hsieh et al., 2015; Rutledge et al., 2015).

What distinguishes a coalescing from a non-coalescing gene, therefore, is not whether it is coordinately regulated, transcribed at a high level, or induced by heat shock. What dictates coalescence is whether a gene is regulated by Hsf1. Indeed, using a combination of conditional Hsf1 nuclear depletion, ectopic Hsf1 targeting and genome editing, we have demonstrated that Hsf1 is both necessary and sufficient to drive the interaction of a transcriptionally competent Pol II gene with other Hsf1-regulated genes. Particularly compelling are two complementary observations: (i) *de novo* interaction of an otherwise non-heat-shock-responsive gene converted to thermal responsiveness and spatial reorganization by ectopic targeting of Hsf1; and (ii) abrogation of interactions between *hsp12-∆HSE* and other *HSP* genes, despite continued robust transcription of *hsp12-∆HSE* in response to heat shock (paralleling similar observations of *HSP12^+^* in cells subjected to Hsf1 nuclear depletion).

It is worth emphasizing that intergenic interactions between Hsf1-regulated genes go well beyond classic transvection phenomena (enhancer - promoter interactions). Indeed, the discovery of coding region interactions between Hsf1-regulated genes, but not of other actively transcribed genes, constitutes a key novel finding of this study. We are unaware of other yeast activators that possess comparable activity, although our experiments do not rule out their existence. The closest example may be that of an erythroid-specific transcription factor, Klf1. Using a combination of 3C, ChIP-3C, FISH and immunofluorescence, Fraser and colleagues have shown that in mouse erythroid cells, Klf1-regulated globin genes relocate into transcription factories where they engage in preferential (although not exclusive) interchromosomal associations with other Klf1-regulated genes (Schoenfelder et al., 2010). Thus, Klf1 drives preferential physical interactions between its target genes in response to a developmental signal; evidence reported here indicates that Hsf1 drives preferential interactions between its target genes in response to an environmental signal. Hi-C and/or ChIA-PET analysis will be required to show whether such interactions are exclusive to the Hsf1 regulon.

### Relationship to Other Examples of Gene Clustering and Repositioning of Active Genes to the Nuclear Periphery

As alluded above, a particularly striking aspect of our study is that constitutively active genes *FRA1* and *PAU17*, despite being located in close linear proximity to *HSP* genes, do not interact with them. Such specificity contrasts with a recent report of methionine-responsive genes in yeast that engage in intrachromosomal clustering upon their induction as assessed by 3C, yet unlike what we observed here, unrelated neighboring genes also tended to interact (Du et al., 2017). More similar to the specificity and selectivity of Hsf1-target gene interactions are observations that TNFα-responsive genes in human endothelial cells engage in intrachromosomal interactions upon cytokine stimulation (Papantonis et al., 2010), whereas an actively transcribed gene interposed between them, and located nearby to one of them, is excluded from such colocalization (Fanucchi et al., 2013).

In addition, microscopy and biochemical analyses have shown that yeast *GAL* genes relocate to the nuclear pore complex (NPC) upon galactose induction (Casolari et al., 2004). Such repositioning has been reported to be accompanied by sustained clustering of *GAL* alleles located on homologous chromosomes, initially at the NPC and subsequently in the nucleoplasm, as detected by a microscopy-based analysis (resolution of ~500 nm) (Brickner et al., 2016) (reviewed in (Du and Bai, 2017)). However, no evidence of *GAL1-10* allelic interaction was seen in galactose-induced diploids using either Hi-C (Kim et al., 2017) or wide-field fluorescence localization microscopy (Backlund et al., 2014). Therefore, it is unclear whether repositioning of the *GAL* locus to the NPC or its inter-allelic clustering is related to the robust and intricate physical interactions that we detect between *HSP* genes using 3C, whose resolution is ~1-5 nm (Dekker and Mirny, 2016). In a similar vein, earlier studies on the effect of heat shock on human nuclear substructure reported the existence of “stress bodies.” As these stress bodies appear to be arrays of HSF1 bound to repetitive, heterochromatic DNA sequences that are spatially independent from *HSP* gene transcription (Jolly et al., 1997; Metz et al., 2004), they are unlikely to be related to the concerted coalescence of *HSP* genes reported here.

Evanescent interactions between Hsf1 target genes contrasts with models suggesting that actively transcribed genes relocate into statically assembled substructures (Feuerborn and Cook, 2015; Mitchell and Fraser, 2008). Rather, our observations resemble the dynamic assembly of Pol II clusters in serum-stimulated human cells (Cisse et al., 2013) or the dynamic sorting of immunoglobulin genes residing on different chromosomes into transcription factories during mouse B-cell development (Park et al., 2014).

### Is *HSP* Gene Coalescence an Example of Phase Separation?

Recently, phase separation of multi-molecular assemblies has been suggested as a mechanism for transcriptional control (Cho et al., 2018; Chong et al., 2018; Hnisz et al., 2017; Sabari et al., 2018). We have described observations consistent with the *HSP* regulon undergoing a liquid-liquid phase separation-like process in response to heat shock. In particular, we have observed that genes sharing in common only the identity of the DNA-bound transcription factor coalesce into diffraction-limited foci under activating conditions. While such coalescence accompanies heightened expression of these genes – and Pol II transcription is indeed required for *HSP* gene interactions as we demonstrate here – the intensity of transcription cannot be the only parameter dictating foci formation. RNA measurements in acutely heat-shocked cells reveal that Msn2/4-regulated *CTT1* and *PGM2* are expressed at levels that equal or exceed several Hsf1 targets studied here, including *HSP12*, *UBI4* and *TMA10* (Figure 2, A-C). Yet *CTT1* and *PGM2* do not detectably interact with one another, nor with representative *HSP* genes.

Why then do Hsf1-regulated genes interact with each other, while Msn2/4-regulated genes do not? One possibility is the presence of low complexity domains (LCDs) / intrinsically disordered regions (IDRs) in Hsf1. IDRs have been postulated to contribute to phase separation of membraneless organelles such as stress granules (Protter and Parker, 2016); those in the abundant nuclear protein HP1 appear to contribute to phase separation of constitutive heterochromatin in both insects and mammals (Larson et al., 2017; Strom et al., 2017). While such low complexity, intrinsically disordered structure may contribute to *HSP* gene coalescence, it cannot be the only reason, since most gene-specific transcription factors, including Msn2/4, also possess IDRs. Indeed, Hsf1 DNA binding in the absence of transcription does not trigger genes to interact *in trans* (Figures 4F, 4H).

Hence, Hsf1 may recruit a distinct set and/or quantity of coactivators and Pol II-associated machinery. Mediator is a particularly attractive candidate, as it is abundant at activated *HSP* genes while nearly undetectable at comparably expressed *CTT1* and *RPL* genes (Anandhakumar et al., 2016; Fan et al., 2006; Kim and Gross, 2013). The relative scarcity of Mediator at *CTT1* and *RPL* may reflect its transient association with these genes despite their strong activation (Jeronimo and Robert, 2014; Wong et al., 2014). Stable association of Mediator at Hsf1-regulated genes may be a key to *HSP* gene coalescence and putative phase separation. Interestingly, IDRs are abundant within this coactivator, and are especially enriched within its Tail module (Toth-Petroczy et al., 2008), the surface of Mediator contacted by Hsf1 (Kim and Gross, 2013). Consistent with this idea, Young and colleagues have recently shown that IDRs in Mediator subunit Med1 contribute to its liquid-liquid phase separation *in vitro* and in mouse cells (Sabari et al., 2018). Therefore, Hsf1-Mediator complexes, along with other covalently modified chromatin-associated proteins and/or nascent RNA transcripts that comprise multivalent networks (Hnisz et al., 2017), may act in a concerted fashion to drive *HSP* gene coalescence.

## MATERIALS AND METHODS

### Yeast Strains

The diploid strains, ASK702 (*HSP104-lacO_256_ HSP12-lacO_128_ GFP-LacI*) and IGY101 (*HSP104-lacO_256_ HSP12-lacO_128_ GFP-LacI*), were created by crossing a MATα derivative of DBY255 with ICY33 and DBY646, respectively (haploid strains generously provided by D. G. Brickner and J. H. Brickner, Northwestern University). ASK706 is a derivative of ASK702 bearing *POM34* C-terminally tagged with mCherry. For fluorescence microscopy of fixed cells, strains ASK706 and IGY101 were used. The diploid strain, SCY712 (*SIK1-mRFP POM34-GFP*), was made by crossing ATY1513 with SCY711.

To construct the diploid strain ASK727 (*HSP104-lacO_256_ TMA10- tetO_200_ GFP-LacI TetR-mCherry*), we first excised *KAN-MX* from ASK701 (*MATα HSP104-lacO_256_ SEC63-Myc13::KAN-MX*), creating VPY101. *KAN-MX* was then integrated downstream of *TMA10*, creating ASK721. Plasmid pSR14, containing *tetO_200_::LEU2* (Rohner et al., 2008) (gift of S. Gasser, Friedrich Miescher Institute for Biomedical Research), was linearized using AscI (generates homologous ends to *KAN-MX*). This linearized DNA was integrated at the *TMA10-KAN-MX* locus, creating ASK722. To create a *MATa TetR-mCherry* expressing strain, we crossed YAM1269 (*MATa leu2::TetR-mCherry::hphMX::leu2*; gift of A. MacQueen, Wesleyan University) with W303-1B (*MATα*), sporulated the resultant diploid, and isolated the desired spore (*MATa leu2::TetR-mCherry::hphMX::leu2* lacking other markers present in YAM1269; see Table 1) termed ASK726. ASK726 was then crossed with ASK722 to obtain the diploid ASK727. We note that *TetR-mCherry* contains an N-terminal nuclear localization signal (A. MacQueen, personal communication).

**Table 1.**
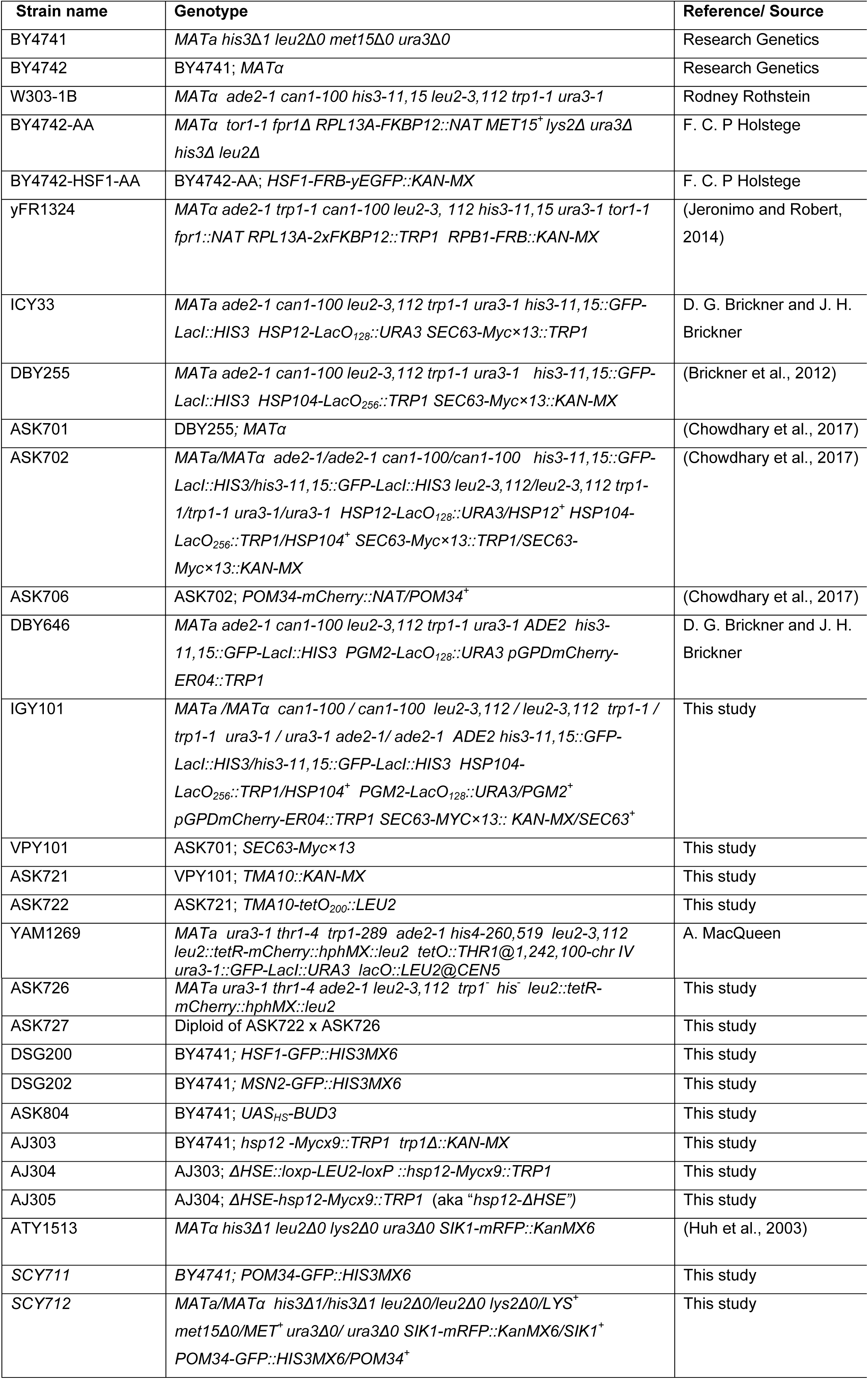
Yeast Strains.

To construct ASK804, a strain bearing the *UAS_HS_-BUD3* transgene, a *loxP-*flanked *KAN-MX* containing fragment was amplified from plasmid pUG6 (Addgene) using primers *BUD3F_loxpF* and *AdaptR_loxpR* (primer sequences are provided in Table S2). Employing BY4741 genomic DNA as template, UAS_HS_ (an 86 bp sequence spanning HSEs 1 − 3 of *HSP82* (Erkine et al., 1999)) was amplified using primers *AdaptF_HSE1F* and *BUD3R_HSE3R*. The two amplicons were then combined in a PCR overlap extension reaction to generate a DNA fragment containing *loxP-kanMX-loxP-3xHSE* flanked by ~40 nt of *BUD3* sequence centered at −182 (with respect to ATG [+1]). This DNA fragment was gel purified and transformed into BY4741; transformants were screened using genomic PCR. The selection marker, *KAN-MX*, was subsequently excised from the desired integrant by ectopic expression of Cre recombinase. The insertion and flanking chromosomal sequence were validated by DNA sequencing (Eurofins Genomics).

To construct AJ305, a strain bearing a deletion of the heat shock element (HSE) upstream of *HSP12*, we used BY4741 *trp1∆::KAN-MX* as the parent. AJ303, a derivative bearing C-terminally tagged *HSP12-Mycx9*, was transformed with a PCR amplicon containing *loxp-LEU2-loxp* in place of the 127 bp DNA sequence spanning −930 → −804 with respect to the ATG pf *HSP12*. The approximate midpoint of this deleted sequence corresponds to the HSE occupied by Hsf1 (see Figure 5A, *right*). The template for overlap extension was pUG73, a plasmid bearing *loxP-*flanked *KlLEU2*. Following transformation, the desired *loxp-LEU2-loxp* integrant (*∆HSE::loxp-LEU2-loxP::hsp12-Mycx9::TRP1*) was identified and named AJ304. AJ304 was then transformed with pSH47, a *URA3-CEN* plasmid containing *GAL1-CRE. LEU2* was excised upon expression of Cre recombinase and pSH47 counterselected using standard procedures. The DNA sequence of the *∆HSE-hsp12-Mycx9::TRP1* locus (referred to as “*hsp12-∆HSE*”) was confirmed and the resultant strain named AJ305.

A complete list of strains is provided in Table 1. PCR primer sequences are provided in Table S2.

### Culture Conditions

For 3C analysis, cells (strain BY4741 except where noted otherwise) were grown at 30°C in YPDA (YPD [yeast extract-peptone-dextrose] supplemented with 0.002% adenine) to a mid-log density (A_600_ = 0.65-0.8). A portion of the culture was maintained at 30°C (control conditions; non-heat-shocked (NHS) sample) while the remainder (heat-shocked (HS) sample) was subjected to an instantaneous 30°C to 39°C thermal upshift for the indicted duration as previously described (Chowdhary et al., 2017).

Conditional depletion of select nuclear proteins was conducted using the Anchor Away system (Haruki et al., 2008). BY4742-HSF1-AA and yFR1324 (Rpb1-AA) were grown in YPDA at 30°C. To anchor-away FRB-tagged Hsf1 and Rpb1 proteins, rapamycin was added to a final concentration of 1 μg/ml to early log cultures (A_600_ = 0.4-0.5). BY4742- HSF1-AA cells were incubated for 90 min in the presence of the drug (except where noted below), while yFR1324 cells were incubated for 60 min. At this point, cells were subjected to an instantaneous heat shock at 39°C for 10 min, and then processed for 3C or ChIP analysis. In the case of RT-qPCR analysis (Figure S3A), a heat shock time course (0, 2.5, 15, 45 min) was employed following rapamycin pre-treatment.

For spot dilution analysis, cells were grown to stationary phase in YPDA. Master suspensions for each strain were prepared by diluting the saturated culture to a uniform cell density (*A*_600_=0.5) and were transferred to a 96-well microtiter plate. These were then serially diluted five-fold and 6 μl were transferred onto a solid YPDA or YPDA + 1 μg/ml rapamycin. Cells were grown at either 30° or 37°C for 2-3 days.

### Chromosome Conformation Capture

TaqI-3C was conducted essentially as previously described (Chowdhary et al., 2017). Briefly, cells were grown, heat-shocked at 39°C (or maintained at 30°C) and crosslinked with 1% formaldehyde. Crosslinked cells were harvested from a 50 ml culture and then subjected to glass bead lysis for two cycles (20 min each) of vortexing at 4°C. A 10% fraction of the crude chromatin lysate was digested with 200 U of Taq I at 60°C for 7 h. Taq I was heat-inactivated (80°C for 20 min) in the presence of SDS (final concentration of 1%). Excess SDS was quenched via addition of Triton-X100 (final concentration of 1%). The digested chromatin fragments were centrifuged, and the pellet was resuspended in 100 μl of 10 mM Tris-HCl (pH 7.5). Proximity ligation was performed with 7x diluted Taq I digested chromatin using 10,000 cohesive end units of Quick T4 DNA ligase at 25°C for 2h. The sample was then digested with RNase (final concentration of 30 ng/μl) at 37°C for 20 min. Proteinase K (final concentration of 70 ng/μl) digestion was performed at 65°C for 12 h in the presence of 0.1% SDS. The 3C DNA template was extracted using phenol-chloroform and precipitated in the presence of glycogen.

Quantitative PCR was performed using anchor primers that were paired with primers abutting Taq I sites lying on the same or another gene, to enable detection of intragenic or intergenic interactions, respectively. Locations of 3C primers used in this study are provided in Figure S1.

A notable feature of our procedure was the use of a 4 bp cutter, Taq I, which recognizes sites located within the UAS/promoter, 5’-end, coding region and 3’UTR/terminator regions of most genes evaluated in this study (see physical maps; Figure S1). In addition, to circumvent typical problems associated with 3C-based techniques, such as the difficulty in normalizing raw contact frequencies of different chromosomal regions, we incorporated multiple controls. The most important of these was to normalize each restriction site tested to percent digestion to account for the potential variation in accessibility of local chromatin structure under each physiological condition or genetic context (see Figure S6 for representative examples). For determining percent digestion efficiency at each Taq I site, the region across the restriction site was amplified using convergent primers (sequences provided in Table S2). This step, which to our knowledge is unique to our procedure, greatly alleviates an under-appreciated problem endemic with all 3C-based procedures.

Additional controls incorporated into our procedure include the following: (i) normalization to purified genomic DNA similarly cleaved and ligated to account for variation in primer pair efficiencies; (ii) normalization to a non-cut region of the genome (internal recovery control) to account for variation in the recovery of 3C templates; (iii) normalization to a no-ligation control to ensure a ligation-dependent 3C signal; (iv) normalization to a no-template control to account for primer dimer background; and (v) use of tandem primers to generate 3C signals, minimizing the possibility of 3C products arising from crosslink-independent ligation. (See ***Reviewer’s Appendix*** for algorithms used to calculate normalized 3C interaction frequencies.)

We note that for genes with closely related paralogues (*HSP82* and *SSA2*, whose coding regions bear 92% and 96% sequence homology to *HSC82* and *SSA1*, respectively), we typically used primers with multiple (and/or 3’-end) mismatches to the paralogous sequence to maximize specificity of the tested interaction. At certain Taq I sites, adequate mismatches within nearby DNA sequence did not exist (e.g., *HSP82* +986, *HSP82* +1838, *SSA2* +855, *SSA2* +1905) and these loci were not evaluated.

### ChIP Followed by High-Throughput Sequencing (ChIP-Seq)

Chromatin was isolated from a 600 ml early log culture of BY4741 cells (either NHS or 5 min heat-shocked) and sonicated for 60 cycles on a high power setting using Bioruptor Plus (Diagenode). 10% of the total chromatin extract was immunoprecipitated using a ChIP-grade Hsf1 polyclonal antibody raised in our laboratory (Erkine et al., 1996). Barcoded libraries were generated using 5 ng of the purified ChIP DNA and a ChIP-seq Sample Prep Kit (NEB). These libraries were then sequenced using an Illumina MiSeq genome sequencer. To generate normalized UCSC tracks we combined the replicate aligned BAM files and called peaks with MACS2 using the -B option. Since these are paired-end libraries (-BAMPE), the fragment size was measured directly from the data and MACS2 reported the pileup aligned reads using the insert size from the paired-end BAM file. As a default, the bedGraphs were not normalized by read depth but each bedGraph coordinate intensity was multiplied by a normalization factor (10 million divided by total fragments in the library).

### RNA-seq

RNA-seq read counts were obtained from ribosomal RNA-depleted total RNA isolated from BY4742-Hsf1-AA cells cultivated at 30°C in the presence or absence of 1 μM rapamycin for 45 min (Hsf1-nuclear depleted or Hsf1-containing cells, respectively) prior to continued incubation at 30°C or heat shock at 39°C for 10 min. HTseq-count was used to generate counts and DEseq2 was used to normalize counts across co-sequenced libraries.

### Reverse Transcription-qPCR (RT-qPCR)

RT-qPCR was performed as previously described (Anandhakumar et al., 2016).

### Fluorescence Microscopy

Fluorescence microscopy of *lacO-*tagged, *tetO*-tagged and *Nop56-RFP*-containing strains (both live-and fixed-cell imaging) was conducted as previously described (Chowdhary et al., 2017), except that for analysis of the diploid strain ASK727 (*HSP104-lacO_256_ TMA10-tetO_200_ GFP-LacI TetR-mCherry*), we used an Olympus UPlanFl 100/1.3-NA objective attached to a Photometrics Prime 95B CMOS camera. All other images were taken using a CoolSNAP HQ Charge coupled device camera. For subnuclear localization, a cell was scored positive for coalescence between two fluorophore-tagged genes if the distance between the centroids of fluorescence was <0.4 μm apart. Nine *z* planes, covering the entire depth of the nuclei, were assessed for location of tagged genomic loci.

For Hsf1-GFP and Msn2-GFP live cell imaging, cells were grown in synthetic dextrose complete (SDC) medium containing 2% glucose and supplemented with 0.1 mg/ml adenine until early log phase. 1 ml of culture was harvested and washed with 1 ml of SDC media. An aliquot of cells were pipetted onto a 3% agarose pad made with SDC medium, covered with a cover slip and then imaged. Samples were subjected to heat shock by heating the objective from room temperature to 38°C using a Bioptechs objective heater system. The 89021 filter set was used (Chroma Technology) for imaging GFP and RFP/mCherry. Images were taken through an Olympus Ach 100/1.25-numeric-aperture (NA) objective using a CoolSNAP HQ charge-coupled-device camera. Slidebook Software, version 4 (Intelligent Imaging Innovations), was used to control camera acquisition and the z-axis stepping motor (Ludl Electronic products). Images were analyzed using ImageJ (1.48v) software.

## AUTHOR CONTRIBUTIONS

S.C., A.S.K. and D.S.G. conceptualized the project. A.S.K., S.C. and D.P. performed the experiments. A.S.K., S.C. and D.S.G. analyzed the data. S.C. and A.S.K. made the figures. D.S.G. wrote the manuscript with assistance from A.S.K., S.C. Funding for this project was acquired by D.S.G., S.C., A.S.K and D.P.

## ACKNOWLEDGEMENTS

We thank Jayamani Anandhakumar and Michael Guertin for experimental and bioinformatics assistance, respectively; Kelly Tatchell for assistance with fluorescence microscopy; Rini Ravindran, Ishita Ghosh and Vickky Pandit for strain construction; and Jason and Donna Brickner, Susan Gasser, Frank Holstege, Amy MacQueen, Anne Cornelis Meinema and François Robert for generous gifts of strains and plasmids. This work was supported by grants from the National Science Foundation (MCB-1025025, MCB-1518345) and the NIH (R15 GM128065) to D.S.G., Ike Muslow predoctoral fellowships to S.C. and A.S.K., and an NIH Early Independence Award (DP5 OD017941) to D.P.

## Supplemental Information

**Chowdhary, Kainth, Pincus & Gross**

**Figure S1.**
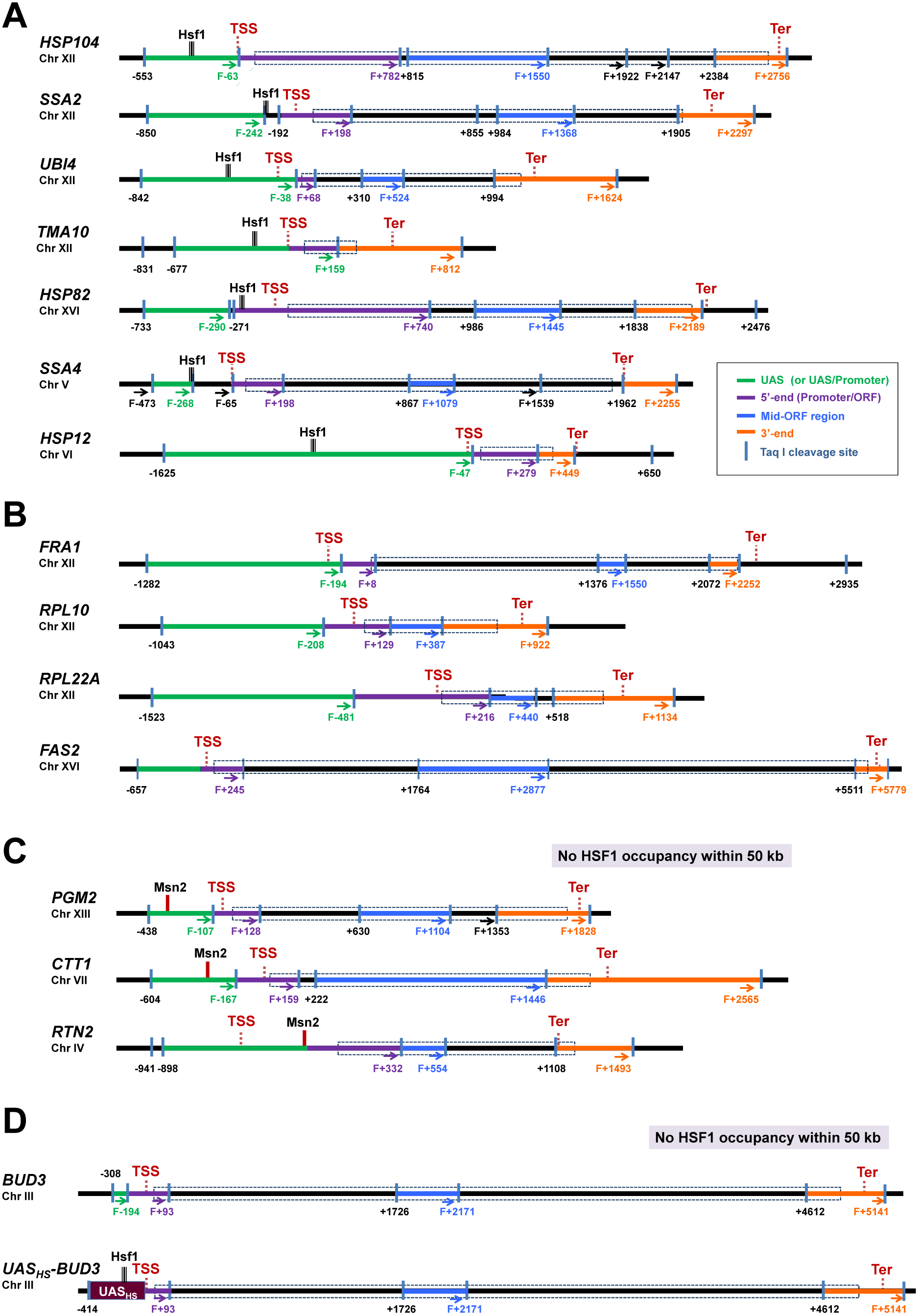
Physical Maps of Genes Evaluated in This Study. (A) Hsf1-regulated genes. Coordinates correspond to Taq I sites (shown as vertical blue bars); numbering is relative to ATG (+1). Primers used for 3C analysis were sense-strand identical (Forward, F) and positioned proximal to Taq I sites as indicated (arrows). UAS/Promoter, 5’-end, Mid-ORF and 3’-end regions are color-coded as indicated. Transcription start sites (TSS) and termination sites (Ter) (Jiang and Pugh, 2009; Xu et al., 2009) are indicated, as are Hsf1 binding sites as determined by ChIP-seq (Figures 1A, 1D, 5A and data not shown). Blue-dotted boxes denote locations of ORFs.
(B) Constitutively expressed genes, as above.
(C) Msn2/Msn4-regulated genes, as above.
(D) *BUD3* and the chromosomal transgene *UAS_HS_ - BUD3*, as above.

**Figure S2.**
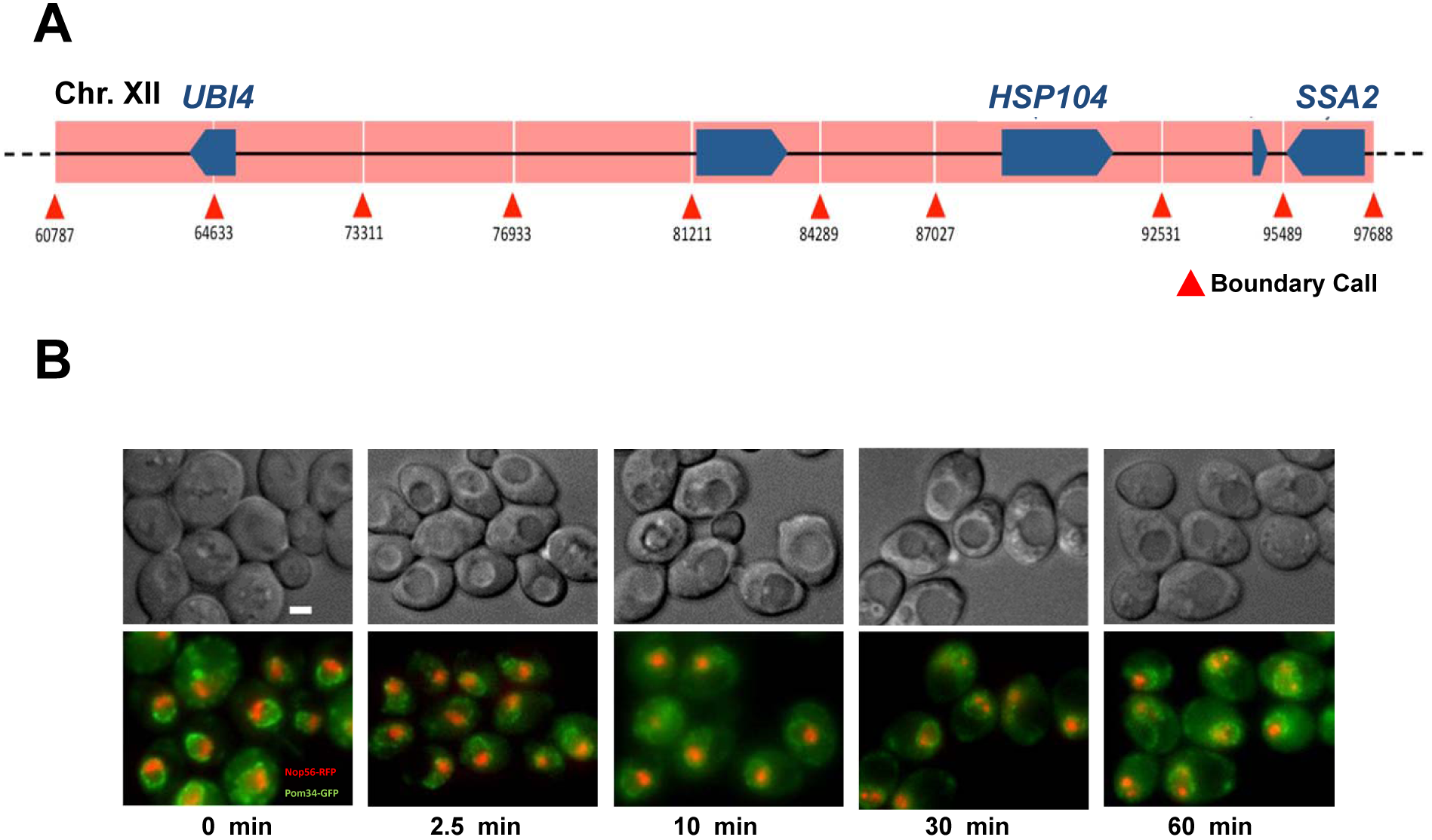
Nucleolar Structural Integrity Is Retained Upon Heat Shock. (A) Chromosomal interaction domain (CID) map of the indicated region within Chr. XII as defined by Micro-C (Hsieh et al., 2015). CID boundaries are indicated.
(B) Fixed-cell microscopy of a diploid strain, SCY712, grown to early log phase at 30°C, then subjected to an instantaneous 39°C heat shock for the times indicated prior to fixation. (*Upper row*) Representative differential interference contrast (DIC) images. (*Lower row*) Corresponding fluorescence images. Red, Nop56-RFP (component of the nucleolus); Green, Pom34-GFP (component of the nuclear pore complex (NPC)). Images were taken across nine *z-*planes with an interplanar distance of 0.5 μm. Shown are maximum intensity projections of planes with above-background nucleolar signal. Note that nucleolar morphology in a projection image may appear slightly different than it does in individual planes, dependent upon nuclear orientation within a given cell. Scale bar, 2 μm.

**Figure S3.**
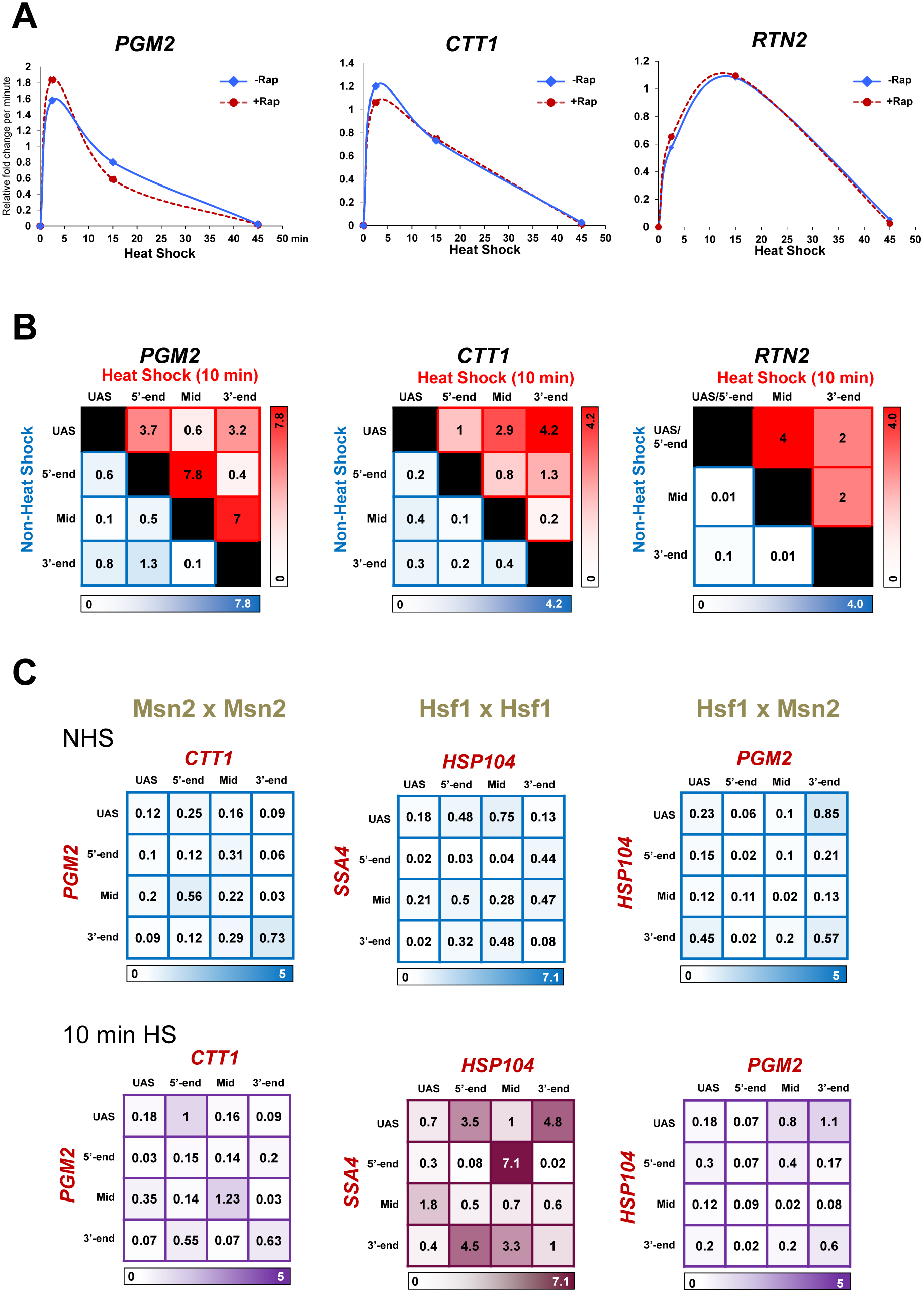
Msn2/4-Target Genes, Despite Strongly Looping Upon Heat Shock, Fail to Engage in Intergenic Interactions. (A) Transcription rates of Msn2/4-regulated *PGM2*, *CTT1* and *RTN2* during a heat shock time course as deduced from steady state mRNA measurements (Reverse Transcription-qPCR). RNA was isolated from Hsf1-AA cells that were pretreated with 1 μg/ml rapamycin for 90 min (red line plot) or not (blue line plot), followed by heat shock for the indicated times. Demonstrates that the increase in *PGM2*, *CTT1* and *RTN2* RNA levels is independent of Hsf1 (see also RNA-seq analysis; Figures 2A and 2B).
(B) Intragenic contact frequencies between the indicated regions of *PGM2*, *CTT1* and *RTN2* under NHS (blue shading) and 10 min HS (red shading) conditions. Intensity of color is proportional to the frequency of interaction. Analysis conducted as in Figure 1B; regions within these genes are defined in Figure S1C. For each pairwise test, N=2, qPCR=4.
(C) Intergenic interaction frequencies between the indicated gene pairs under NHS and 10 min HS conditions (*Upper* and *Lower* matrices, respectively), conducted as in Figure 1B.

**Figure S4.**
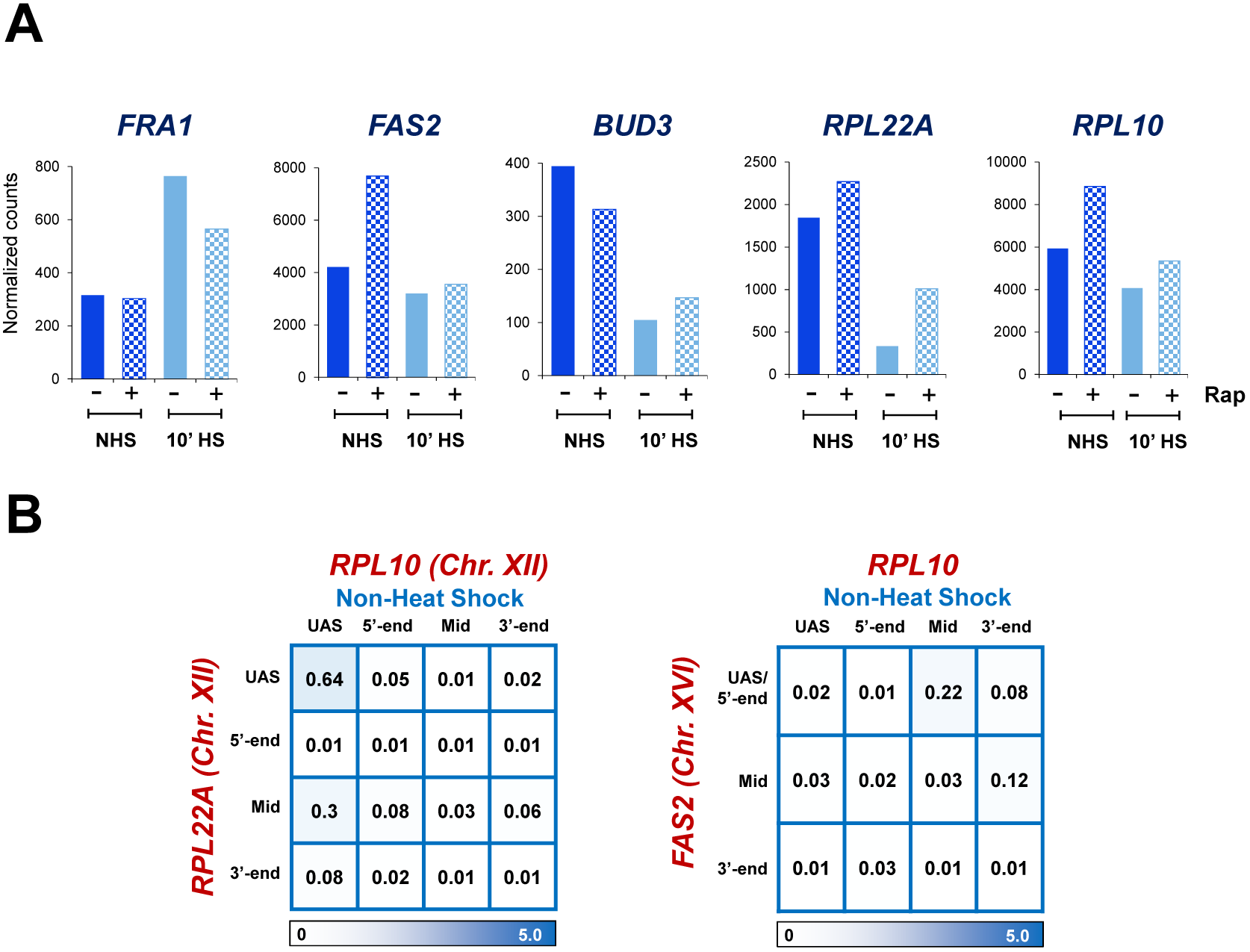
Despite High Levels of Expression, Constitutively Expressed Genes Fail to Engage in Intergenic Interactions. (A) Transcript levels (normalized RNA-seq reads) of select constitutively expressed genes in Hsf1-AA cells. Analysis conducted as in Figure 2, A-C.
(B) Intergenic interactions between constitutively transcribed genes *RPL10, RPL22A* and *FAS2* under NHS conditions. Analysis conducted as in Figure 1B. For each pairwise test, N=2 and qPCR=4. Physical maps are provided in Figure S1B.

**Figure S5.**
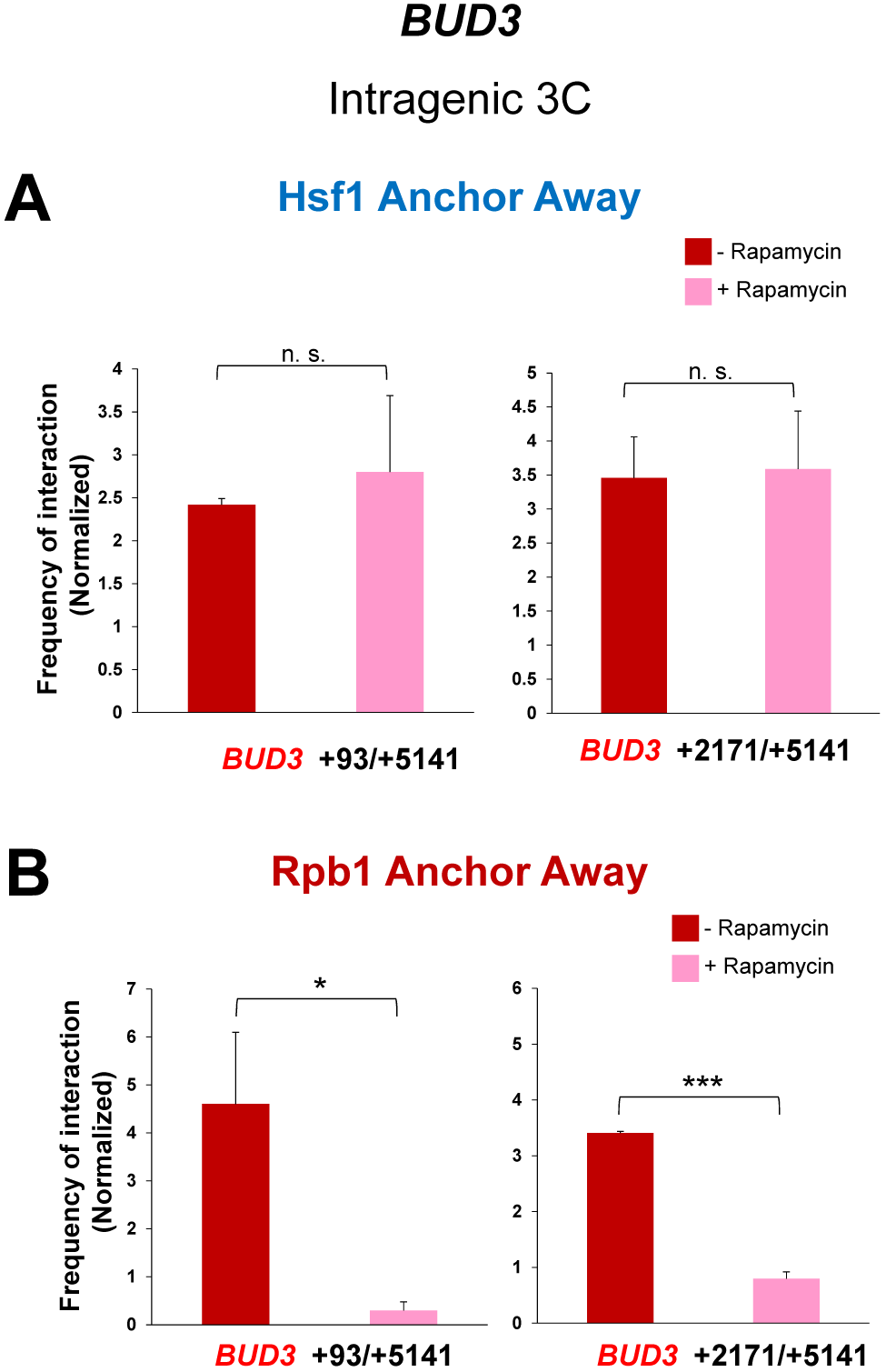
Intragenic Interactions Within *BUD3* Are Suppressed by Nuclear Depletion of Rpb1. (A) Normalized intragenic interaction frequencies within *BUD3* in Hsf1-AA cells pretreated with rapamycin for 90 min or not as indicated, then subjected to a 10 min HS and Taq I −3C analysis. Primer pairs used are indicated below each graph. Demonstrates that intragenic interactions within *BUD3* are independent of Hsf1.
(B) Normalized intragenic interaction frequencies within *BUD3* in Rpb1-AA cells pretreated with rapamycin for 60 min or not as indicated, then subjected to a 10 min HS and Taq I −3C analysis as in A. Demonstrates that intragenic interactions within *BUD3* are dependent on Pol II.

**Figure S6.**
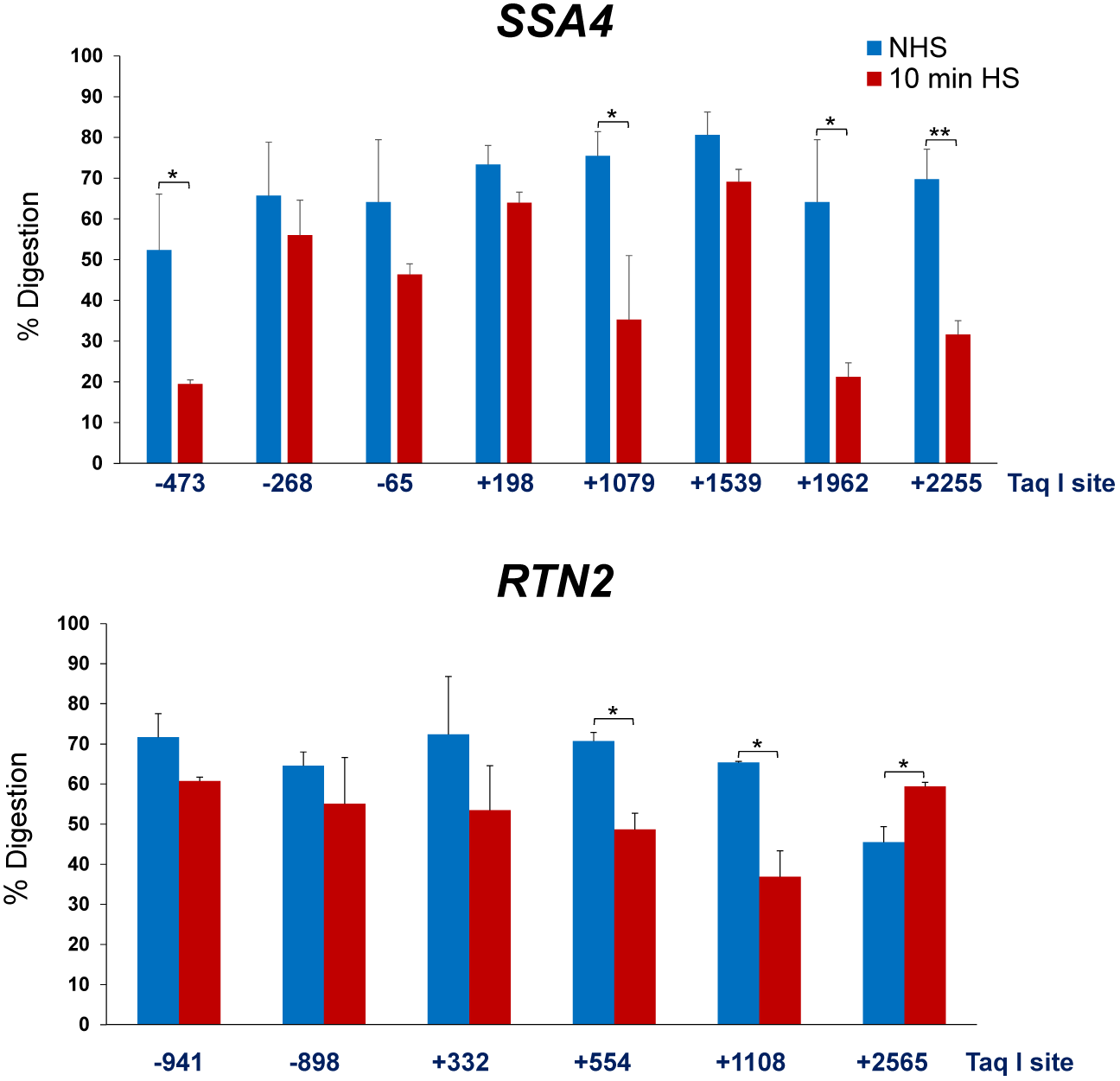
Taq I Digestion Efficiency of Crosslinked Chromatin. Percent digestion of Taq I sites within Hsf1-and Msn2/4-regulated genes under the indicated conditions in BY4741 cells (for additional examples, see Chowdhary et al, 2017). Quantification was done using qPCR as described previously (Chowdhary et al., 2017; Hagege et al., 2007). *, *P*<0.05; **, *P*<0.01 (calculated using one-tail *t*-test). Primers used are provided in Materials and Methods.

**Table S1.** Chromosomal Latitudes of Genes Evaluated in This Study.

| Gene | Chromosome | Location Relative to Centromere (kb) | Location Relative to Telomere on Same Arm (kb) | Exhibit Coalescence? |
| --- | --- | --- | --- | --- |
| **Hsf1-Regulated** |  |  |  |  |
| *HSP12* | VI | 41.0 (L arm) | 107 | **Yes** |
| *HSP82* | XVI | 459.5 (L) | 96.5 | **Yes** |
| *HSP104* | XII | 62.2 (L) | 88.6 | **Yes** |
| *SSA2* | XII | 55.3 (L) | 95.6 | **Yes** |
| *SSA4* | V | 213 (R) | 212 | **Yes** |
| *UBI4* | XII | 86.8 (L) | 64 | **Yes** |
| *TMA10* | XII | >>632 (R) | 295 | **Yes** |
| **Msn2/Msn4-Regulated** |  |  |  |  |
| *CTT1* | VII | 158 (R) | 436 | **No** |
| *PGM2* | XIII | 208 (R) | 449 | **No** |
| *RTN2* | IV | 355 (L) | 94.6 | **No** |

**Table S2.**
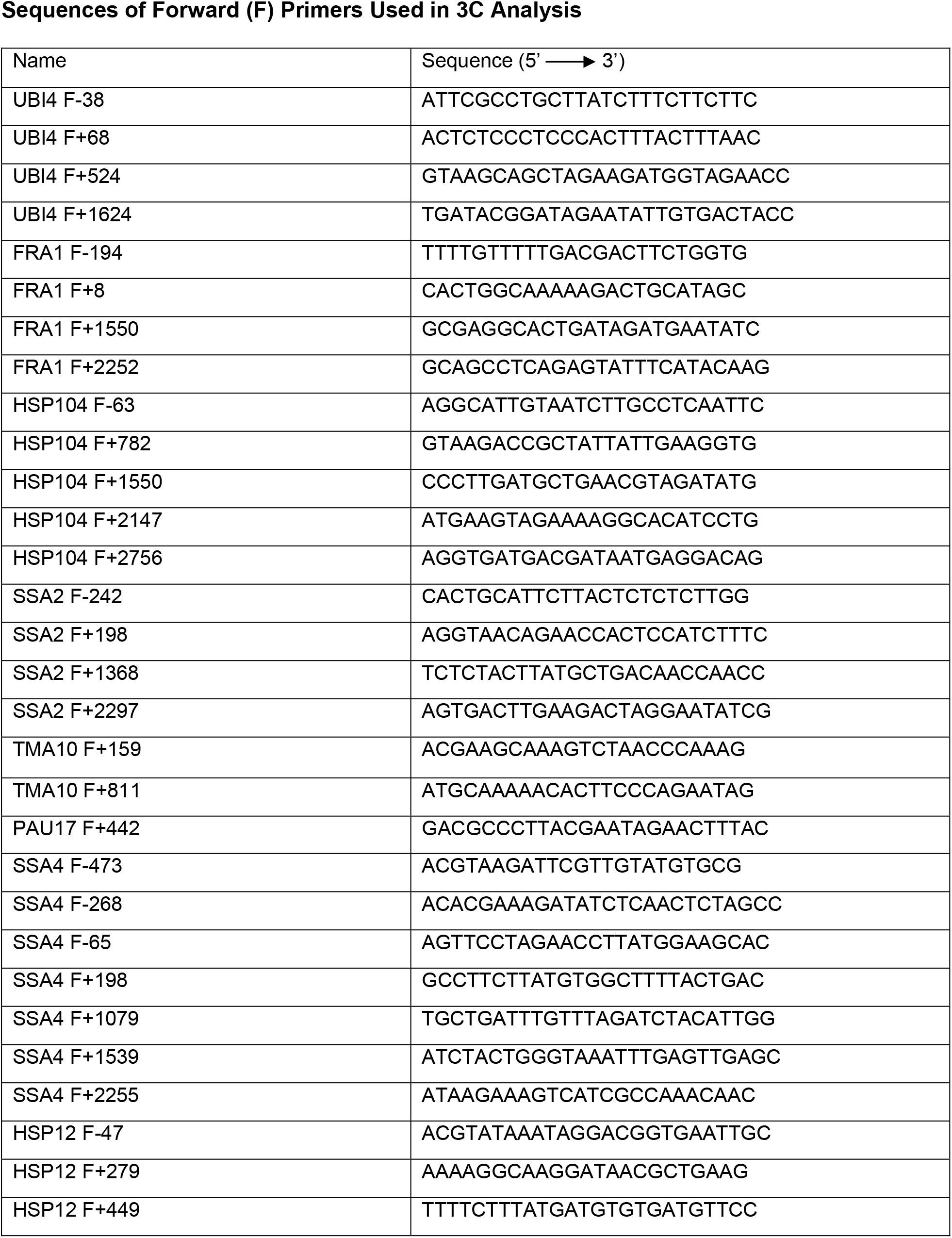
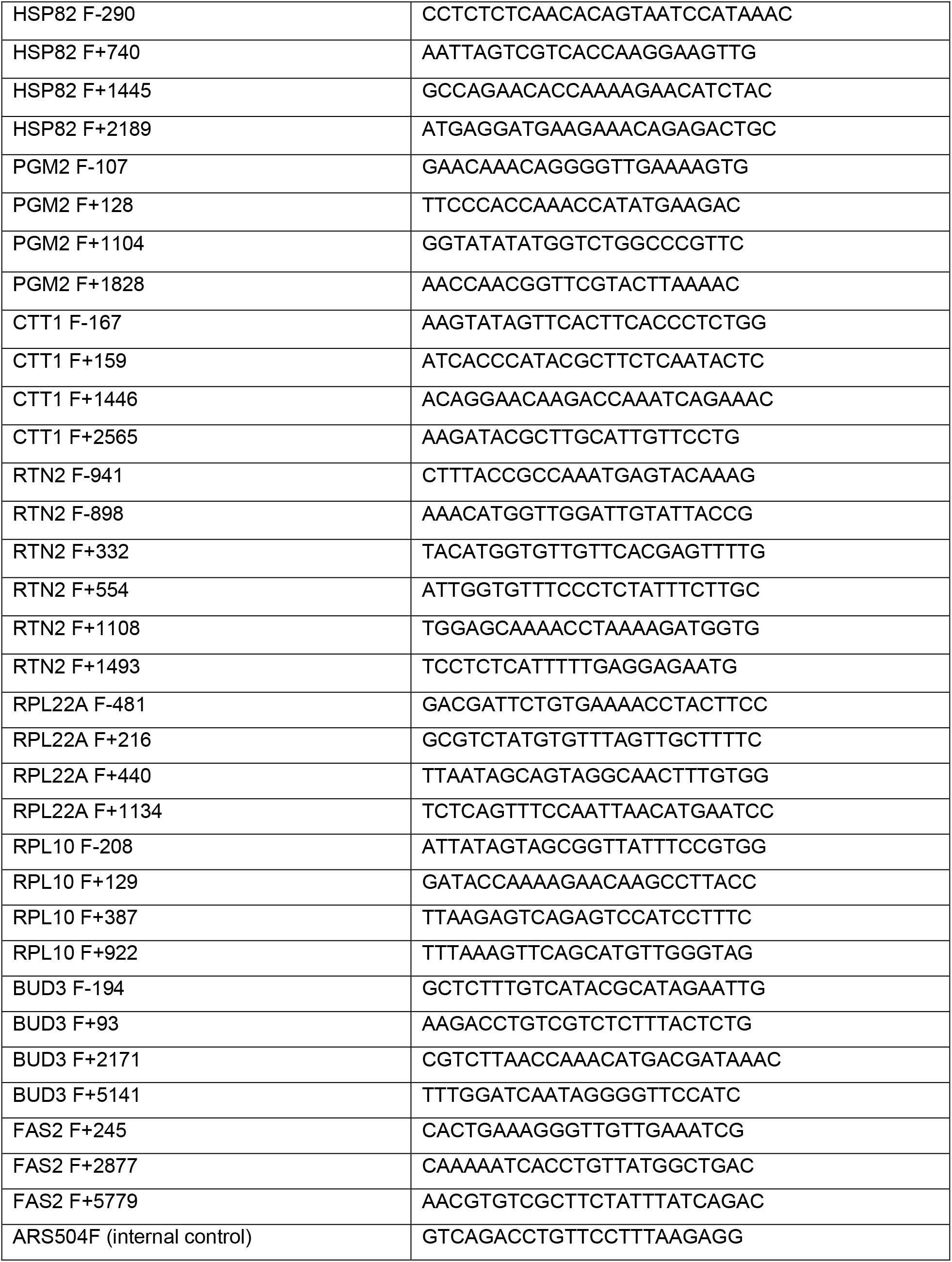
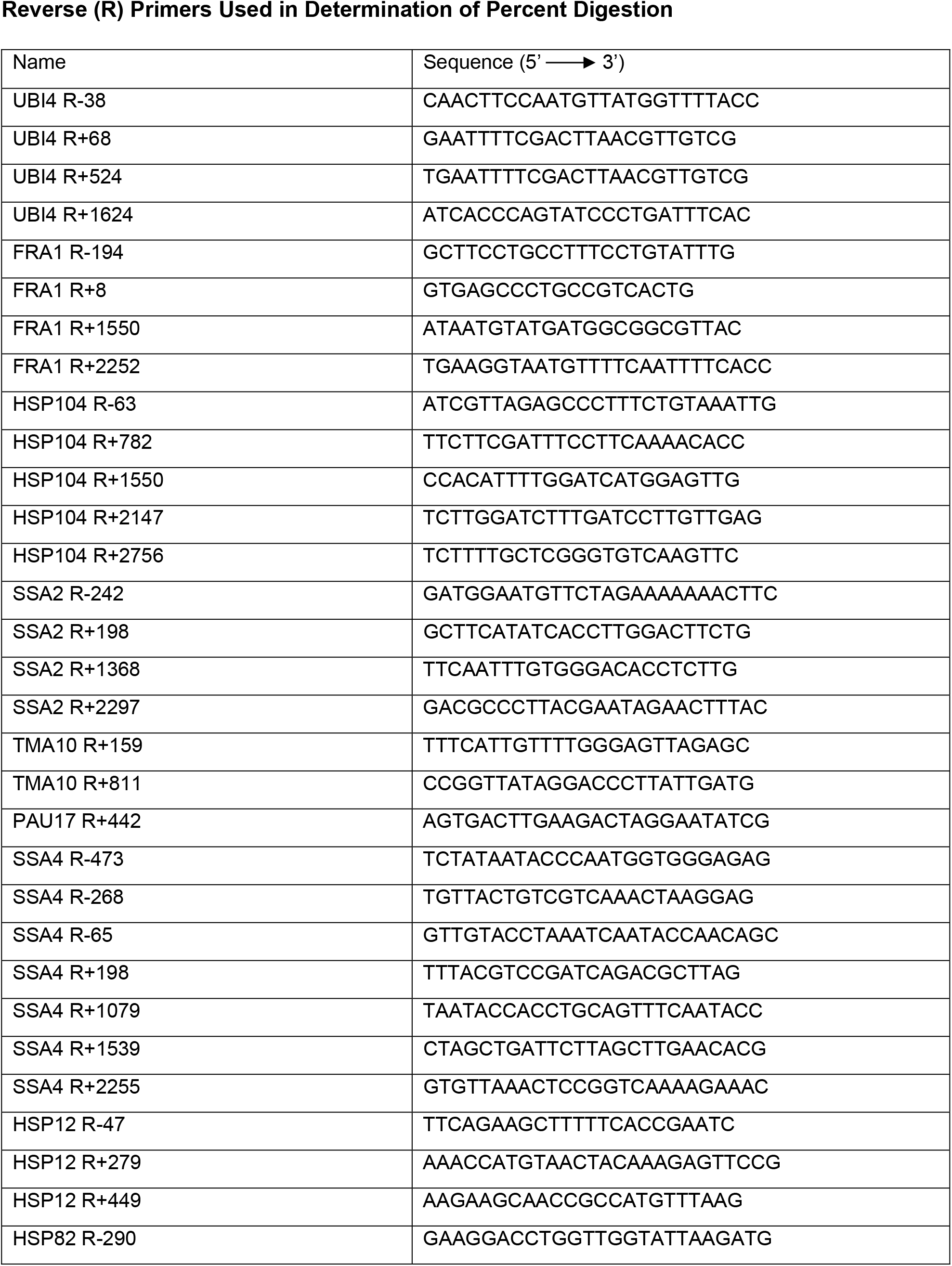
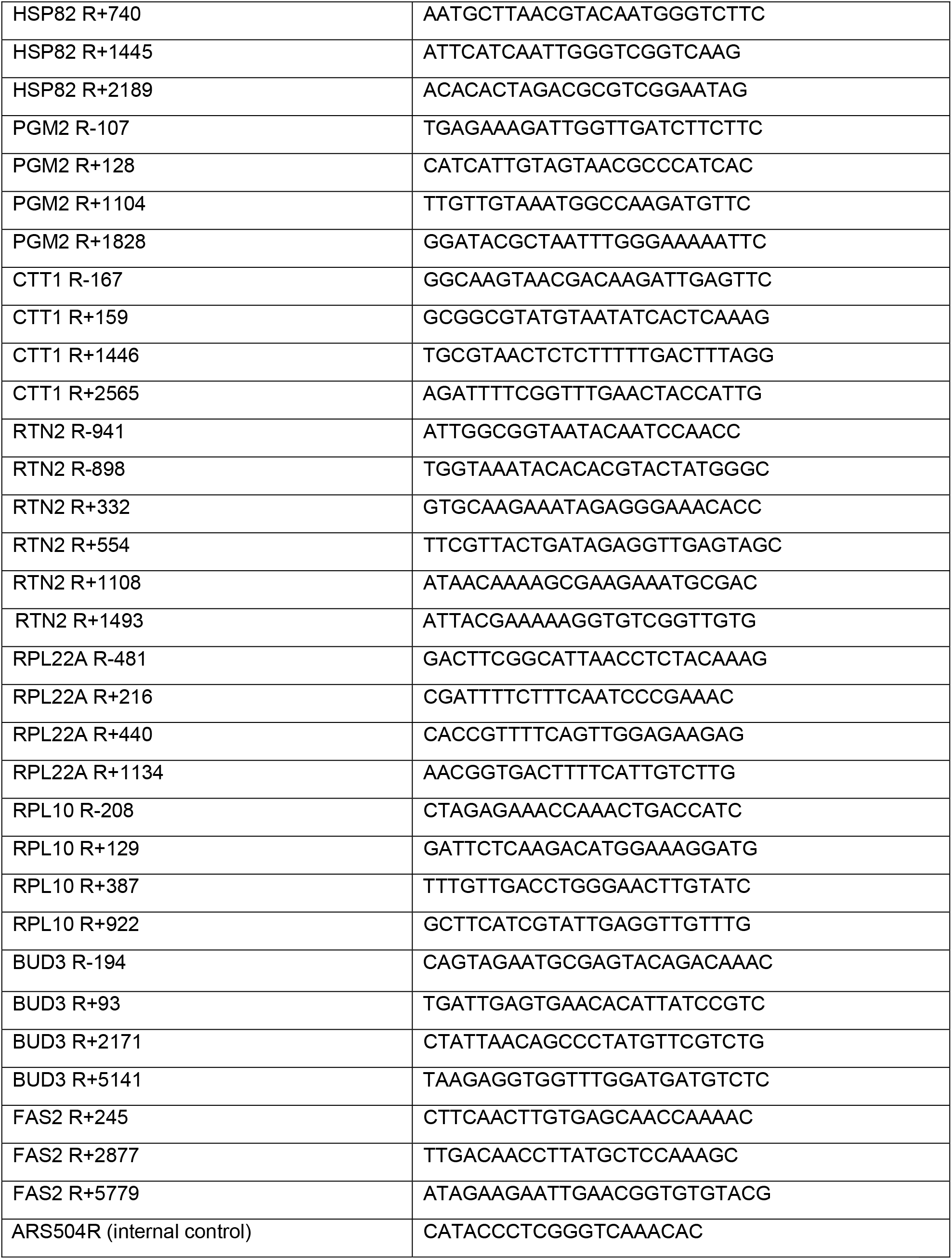
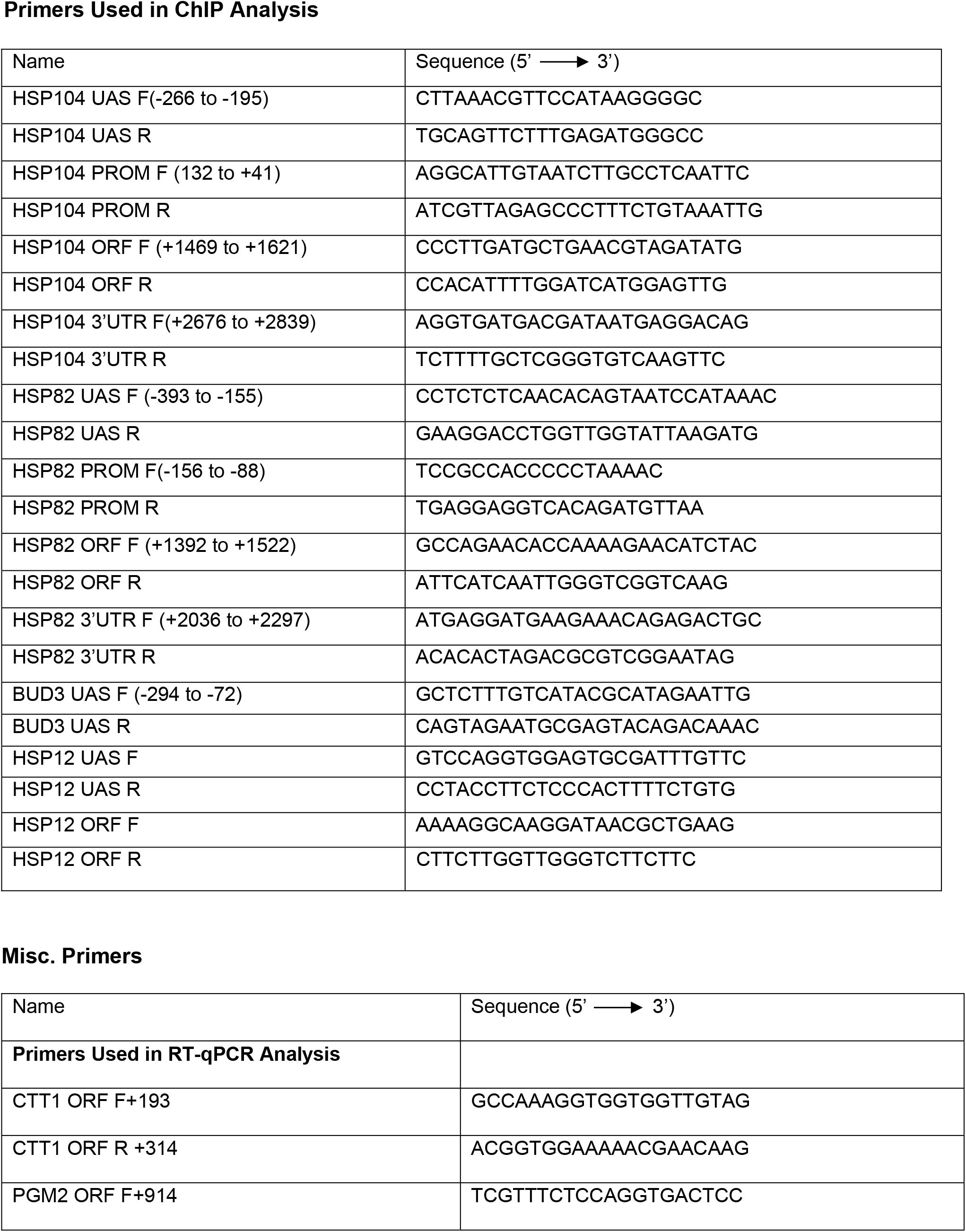
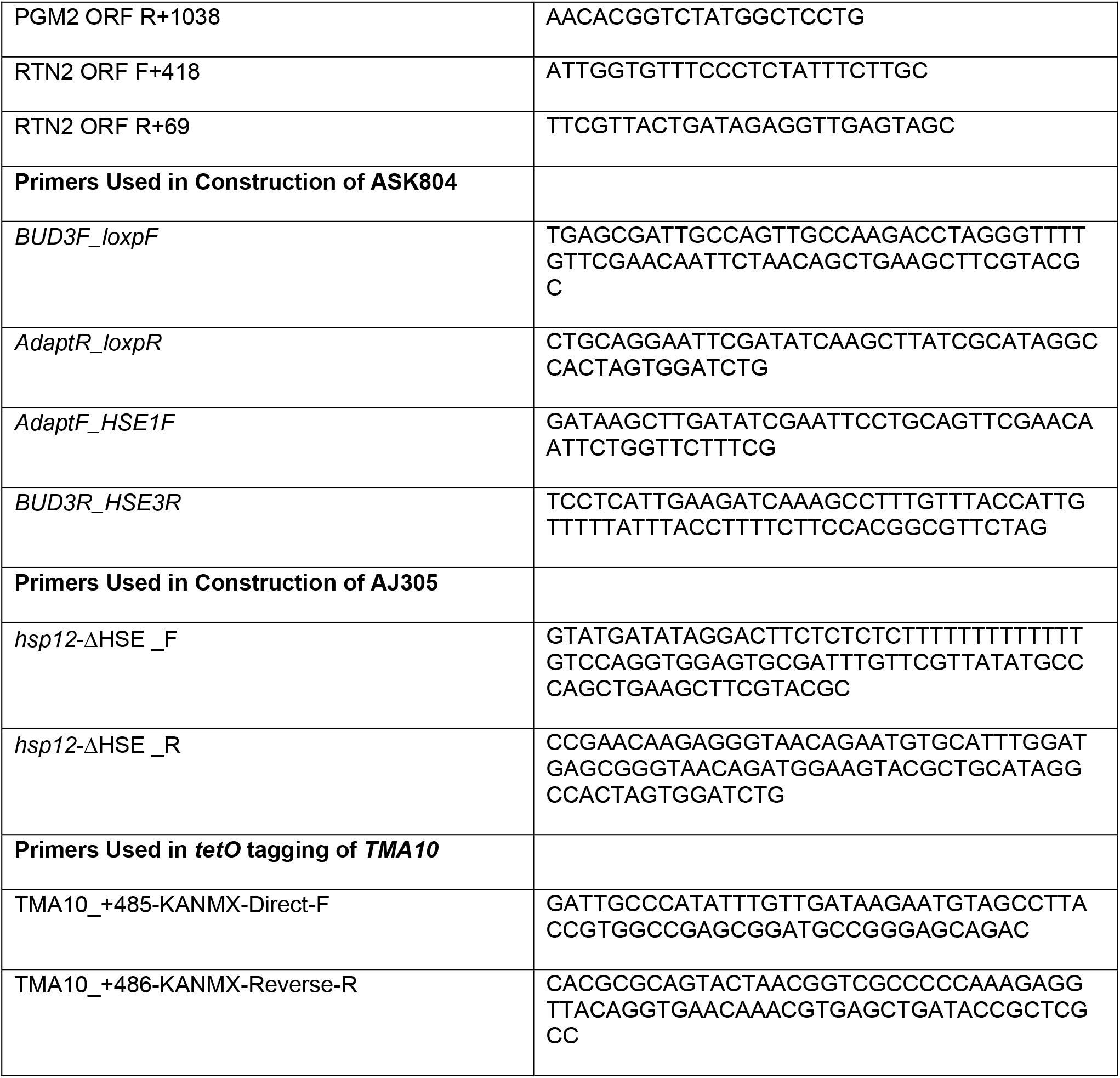
Primers Used in This Study.

